# Synchrotron Tomography-Based Inversion Pipeline for Estimating Elastic Properties of Rat Vertebral Endplate Finite Element Models

**DOI:** 10.1101/2024.09.03.610954

**Authors:** Jishizhan Chen, Alissa Parmenter, Aikta Sharma, Elis Newham, Eral Bele, Sebastian Marussi, Andrew A Pitsillides, Nick J Terrill, Christopher Mitchell, Himadri S Gupta, Peter D Lee

**Affiliations:** Mechanical Engineering, University College London, Torrington Place, London WC1E 7JE, UK; Research Complex at Harwell, Rutherford Appleton Laboratory, Oxfordshire OX11 0FA, UK; School of Engineering and Materials Sciences, Queen Mary University of London, London E1 4NS, UK; Department of Veterinary Basic Sciences, Royal Veterinary College, Royal College Street, London NW1 0TU, UK; Diamond Light Source, Didcot OX11 0DE, UK; Medical, Molecular and Forensic Sciences, Murdoch University, Murdoch 6150, Australia

## Abstract

Lower back pain is linked to vertebral biomechanics, with vertebral endplates (VEPs) playing a key role. Finite element modelling (FEM) is a powerful tool for studying VEP biomechanics but relies on accurate material property inputs, which remain difficult to obtain. Synchrotron computed tomography (sCT) allows for detailed visualisation of the microstructure of intact VEPs under near-physiological loads and, when coupled with digital volume correlation (DVC), can be used to quantify three-dimensional (3D) strain fields, providing experimental reference data for FEM validation. We developed an inversion pipeline to spatially couple DVC data with an image-based FE model, and thus to estimate the elastic properties of rat VEPs. On the first rat lumbar FEM, the pipeline estimated a VEP elastic modulus of 129 MPa and a Poisson’s ratio of 0.24. Welch’s ANOVA revealed statistically significant differences between FEM and DVC strain distributions (p < 0.001) but with small effect sizes (η² = 0.1%–0.6%), indicating high practical similarity. Its efficacy was further validated using Bland–Altman analysis, demonstrating over 95% spatial agreement between the FEM-predicted strains and the DVC measurements across multiple loading steps. The pipeline’s consistency was further evaluated across multiple rat lumbar FE models (n = 3), yielding an estimated VEP elastic modulus = 145 ± 18 MPa and a Poisson’s ratio = 0.28 ± 0.04. Statistically significant regional variations of strain distribution in VEPs were also identified (p < 0.05 to p < 0.001, η² up to 41.8%). This study highlighted the efficacy of the developed pipeline in estimating the isotropic elastic modulus and Poisson’s ratio of VEP FEMs in a physiologically relevant, complex load transfer system. Our pipeline may be used in estimating properties of VEP in larger animals and humans.

## Introduction

Lower back pain is common in all countries and age groups[1], and is the leading worldwide cause of disability[2], particularly in the aged[3]. The exact cause of most cases of lower back pain remains unknown, and due to the limited effectiveness and potential side effects of invasive treatments, current management primarily aims to relieve pain[4–5] rather than addressing underlying biomechanical factors. Recent evidence has identified links between lower back pain and intervertebral disc degeneration, particularly those associated with structural and mechanical changes in the vertebral endplate (VEP)[6–8]. Model-based approaches like finite element modelling (FEM) are a powerful tool for investigating biomechanics in orthopaedics. However, functional modelling at the organ level remains a significant challenge due to its atypical morphology[9–10], heterogeneous composition, and complex loading conditions[11].

To enable accurate VEP modelling, synchrotron computed tomography (sCT) has emerged as a synergistic tool. sCT has recently enabled the visualisation of microstructures under static conditions in intact bovine tails[12], equine digit[13], rat temporomandibular joint[14], mouse knee joints[15–16], as well as human knee joint[17] and foot and ankle[18]. sCT has also proved effective in microstructure evaluation under compressive load in bovine cartilage and meniscus[19], rat lumbar[20–21], and mouse knee joints[20, 22]. The method of sCT allows greater resolution, faster imaging speed, and improved contrast in untreated samples, compared to lab-based CT. The images generated from sCT also enable the application of digital volume correlation (DVC) to quantify three-dimensional (3D) displacement and strain fields in response to uniaxial compression[20–24], similarly to the combined lab CT/DVC approach. This combined sCT/DVC approach provides experimental reference data to validate and increase the accuracy FE models, which is essential for extending simulations to a wider range of orthopaedic conditions, such as multi-axis motion and joint pathophysiology. However, the accuracy of FEM simulations relies on the quality of the material properties and boundary condition inputs. While human spine properties are well-studied, measurement challenges, such as limitations of *in vivo* measurement, individual variability, and reginal heterogeneity, complicate their direct application in individualised FE models. For animal models, such as rats, accurate inputs for spine material properties are equally lacking. These facts render development of a method for their calibration critical to an effective FEM and robust predictions of underlying VEP biomechanics.

CT-based DVC coupled with FEM has been used for comparison of FEM predictions with DVC measurements of strain and displacement[25–28] and has recently been applied as an inverse method for estimating the Young’s modulus of materials, such as human isolated single femoral trabeculae[29]. However, prior property inversion studies have relied on sub-samples and single materials in a simple system, lacking accuracy at the level of spatially paired nodes and validation in a physiologically relevant complex load transfer system (such as intact joints). Moreover, existing inversion approaches exclude the optimisation of Poisson’s ratio, which can lead to significant inaccuracy[30]. To address these challenges, this study aims to develop an inversion pipeline using experimental DVC data as references to simultaneously estimate the effective elastic modulus and Poisson’s ratio of VEP in intact rat lumbar segments. The developed pipeline lays the foundation for future translational applications in both large animal models and human spine biomechanics.

## 2 Results

### 2.1 Establishment of a rat lumbar segment finite element model

The anatomic boundary of VEPs, cartilaginous endplates (CEPs), vertebrae, growth plates, annulus fibrosus and nucleus pulposus can be clearly identified in the sCT image (**Figure 1a** and **1b**). For the VEP, the sCT revealed multiple protrusions (which are significant in rodents but not in humans) (**Figure 1b** and **1e**) within the VEP macrostructure and identified 25-30 annulus fibrosus layers depending on location.

**Figure 1.**
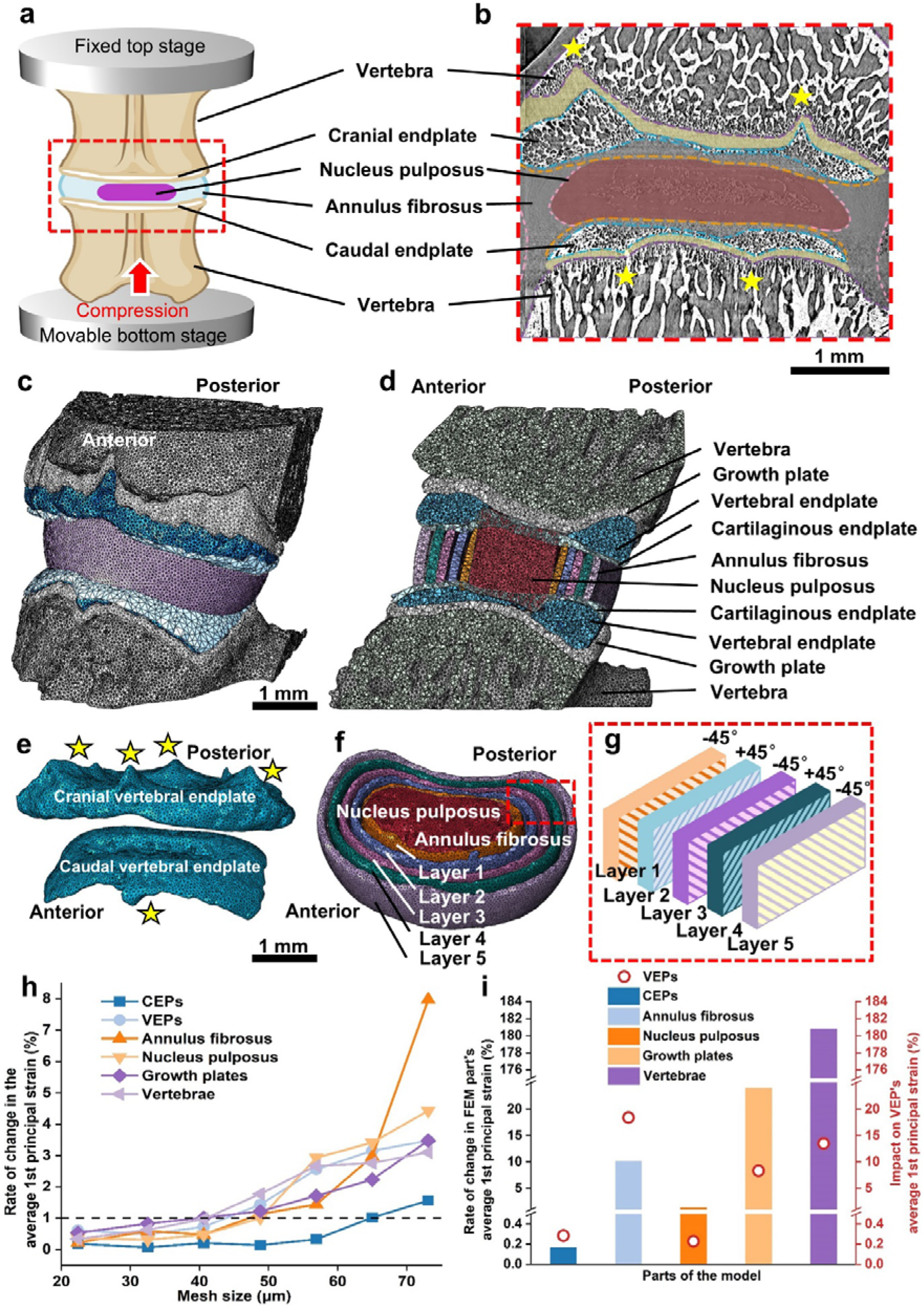
FEM and caudal VEP grid convergence study. **(a)** Experimental setup. Rat lumbar segments were mounted onto the Deben CT500 stage. The red dashed line indicates the scan area. **(b)** Grayscale sCT ortho-slice of vertebrae. Dashed lines and colour blocks indicate the anatomical boundary of the VEPs (blue), CEPs (orange), vertebrae (purple), growth plates (yellow), annulus fibrosus (pink), and nucleus pulposus (dark red). Stars indicate the identified protrusions; **(c)** Integrated FEM of rat lumbar segments generated from (b), showing **(d)** cross-sectional view; **(e)** Isolated FEM of VEPs. Stars indicate the identified protrusions; **(f)** Isolated FEM of nucleus pulposus and multilayer annulus fibrosus. The red dashed line box shows **(g)** the alternating ± 45° anisotropic material orientation of each layer; **(h)** Mesh convergence study on all parts showing the correlation between the average first principal strains and the mesh size. The horizontal dashed line at 1 % represents the level being considered as convergence; **(i)** Sensitivity analysis on all parts. The bar charts and the left *y* axis indicate the impact of extreme modulus values on the parts themselves, whilst the red dots and the right y axis stand for the resulting impact on the VEPs.

The sCT data was used to define the geometry of the FE model. The protrusions were fully resolved in the model, but the annulus fibrosus layers were simplified to a five-layer lamellar structure (each thickened by a factor of five, see **Figure 1c, 1d,** and **1f**) with an alternating ± 45° orientation homogeneous within each layer (**Figure 1g**). To ensure mesh density would not affect FEM predictions, a mesh-convergence analysis was conducted on all parts (VEPs, CEPs, vertebrae, growth plates, annulus fibrosus, and nucleus pulposus). Overall, it was found that average first principal strains diminished steeply (by 3%–68%) along with the decreasing mesh size, before attaining a plateau of less than 1% diminishing rate (**Figure 1h**). Specifically, convergence was considered at a mesh size = 32.5 μm for VEPs, annulus fibrosus, nucleus pulposus, and vertebrae; a mesh size = 24.4 μm for growth plates; and a mesh size = 56.9 μm for CEPs. The mesh convergence test resulted in a lumbar final FEM containing a total number of 1,338,170 elements, creating an FEM platform for simulations.

Using the converged FE model, a sensitivity analysis evaluated the impact of the material properties of various parts on VEP strain distribution. The results demonstte that extreme modulus values, applied to both adjacent and non-adjacent parts, influence VEP strain to varying degrees (**Figure 1i**). Among all FEM parts, the CEP and annulus fibrosus elastic moduli have the most significant impact on the average first principal strain of the VEP than on themselves. Moreover, the annulus fibrosus contributes to the highest VEP strain deviation of 18%. The growth plate and vertebra contribute to 8% and 14%, respectively, whilst the nucleus pulposus has a minimal effect of 0.2%.

### 2.2 Spatial registration of DVC correlation point clouds to FEM nodes

The result of spatial registration shows that 91.7% of FEM nodes were within the DVC point domains, allowing use of a natural neighbour interpolation method to map DVC data onto FEM nodes. The remaining 8.3% of FEM nodes were mapped using extrapolation by the nearest nodes. As a result of spatial registration, the DVC point cloud was down-sampled from 23,529 to 16,250 points (**Figure 2a**). This generated a new DVC point cloud with identical node numbers and spatial locations as the FE model. A visual inspection revealed that the spatial registration had smoothed high strain values in some regions while creating new high strain values in other areas. To quantify the comparability of the DVC point cloud before and after registration, several statistical analyses were performed.

**Figure 2.**
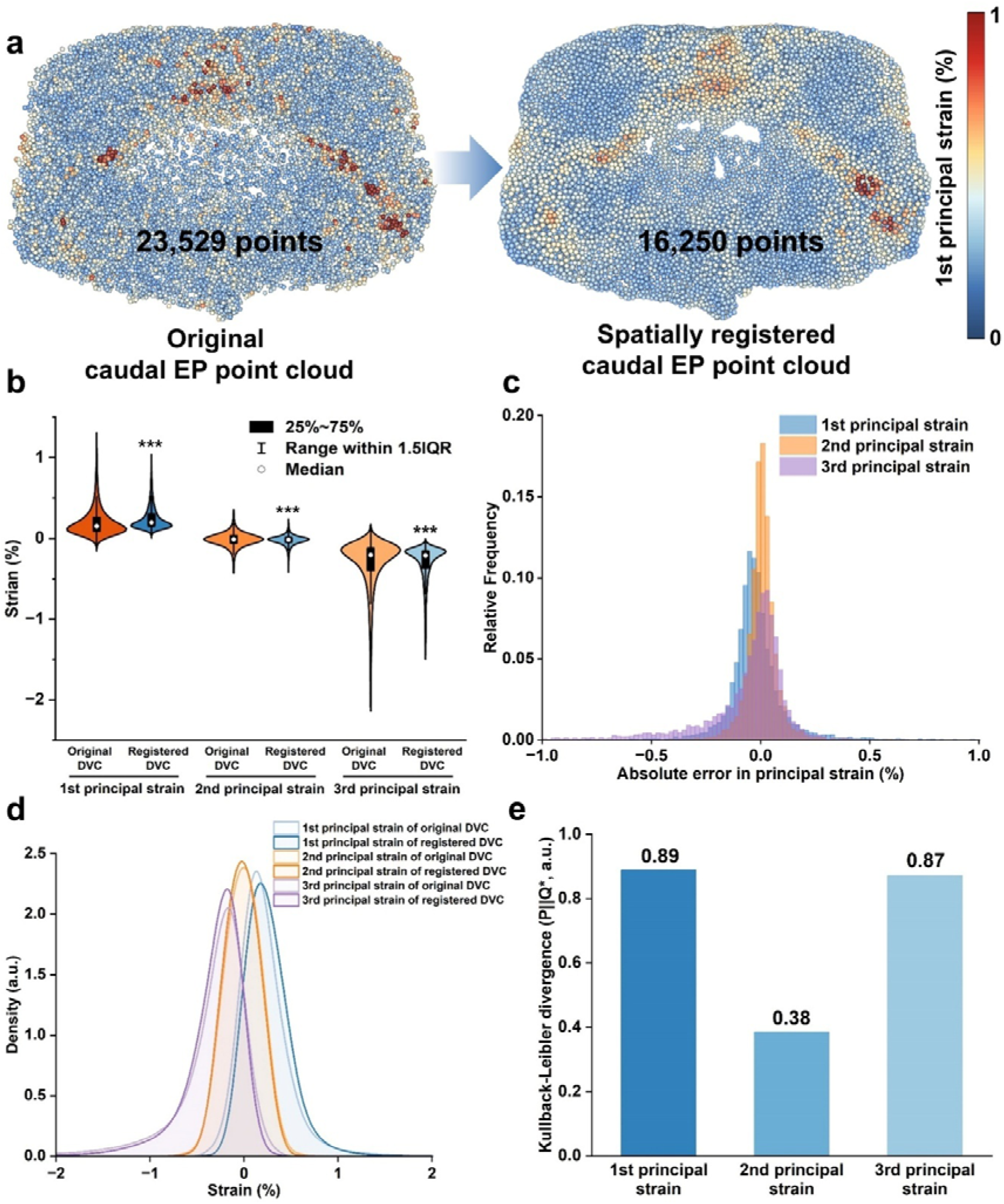
Spatial registration of the DVC point cloud to FEM reference nodes. **(a)** Spatial registration of the DVC point cloud and visualisation of first principal strain; Statistical analyses for deviation evaluation using **(b)** violin plots of strain distribution, **(c)** histogram of absolute error at sub-region level, **(d)** probability density functions at single point level, and **(e)** the corresponding Kullback-Leibler divergence of all principal strains. Not significant (ns): P□>□0.05; *: P□<□0.05; **: P□<□0.01; ***: P□<□0.001; Compared to the original DVC point cloud.

In the strain distribution analysis (**Figure 2b**), smoothing has reduced high strain values, resulting in fewer extremes in the first, second, and third principal strains and a more centralised distribution trend (P < 0.001, η² = 0.2%–0.5%). The median of first, second, and third principal strains increased from 0.15% to 0.19%; from -0.012% to -0.017%; and from - 0.20% to -0.21%, respectively.

The sub-region-based error analysis (**Figure 2c**) shows that, at a voxel resolution of 32.5 μm, the spatial registration causes minor deviation. The largest deviation was observed in third principal strain, with a mean of -0.09%. The histogram of all principal strains has strong peaks near 0% but broad tails, especially for first and third principal strains. The second principal strain is the least affected and displays the smallest spread (0.0048 ± 0.069%), whilst third principal strain is the most affected and shows the largest spread (-0.088 ± 0.32%).

The probability density function (**Figure 2d**) presents the similarity between the original and registered DVC strain distributions at the resolution of single points. The distribution of all principal strains remains highly consistent with minor shift, with the least peak shift in second principal strain from -0.011% to -0.020% and the greatest peak shift in first principal strain from 0.16% to 0.19%. The Kullback-Leibler divergence further quantifies these differences, where first and third principal strains yield high values of 0.89 and 0.87, respectively, indicating slightly greater deviation. The second principal strain has the lowest value (0.38), confirming that its distribution remains the most similar during the registration. The results of Kullback-Leibler divergence are consistent with the shifts in the probability density function (**Figure 2e**).

### 2.3 Inversion pipeline for adjustment of VEP material properties

The inversion pipeline was employed to determine the global optimal VEP material properties, ensuring the best fit between first principal strain values of the caudal VEP in the FEM and those from the DVC. By varying the elastic modulus/Poisson’s ratio combinations we generated 129 local minimum root-mean-square errors (RMSEs) and a global minimum RMSE (2.09 × 10^-3^) corresponding to an elastic modulus E = 129 MPa and Poisson’s ratio ν = 0.24 combination (Figure 3a and **3b**).

**Figure 3.**
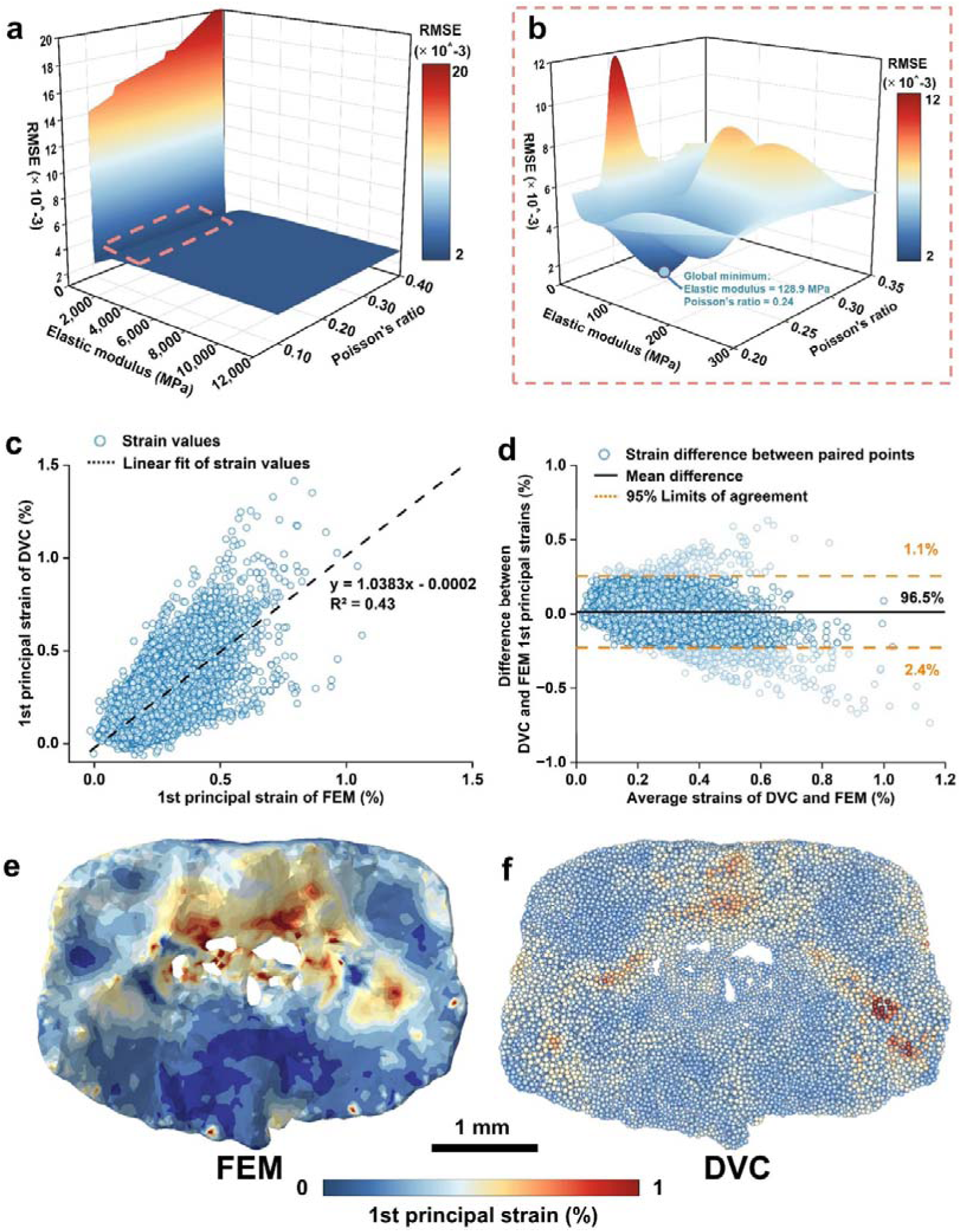
Global optimisation of material properties of caudal VEP and statistical analyses. **(a)** Visualisation of the correlation between first principal strain RMSE and combinations of the elastic modulus and Poisson’s ratio. The pink dashed box indicates **(b)** the zoom-in view of the location of the global minimum RMSE; **(c)** The regression analysis showing the correlation of paired points between FEM and DVC first principal strain. Each dot represents a paired data point; **(d)** The Bland–Altman plot showing the deviations of first principal strain between FEM predictions and DVC measurements. Each dot represents a paired data point; Visualisation of first principal strain on the **(e)** FEM and **(f)** DVC point cloud.

Comparability of this optimised FEM and DVC point cloud reference for the caudal VEP was verified. At paired point level, regression analysis shows a correlation of R^2^ = 0.43 (Figure 3c). In comparison, the Bland–Altman plot[31–32] analyses the global agreement between the two different assays. The result exhibited 96.5% agreement (15,681 nodes, Figure 3d) with poorer agreement only at higher average strain (569 nodes, Figure 3d). FEM visualisation of first and third principal strains demonstrated the greatest deformation in the posterior compartment of the caudal VEP; notably in the protrusions on the vertebral side and consistent with the DVC dataset (Figure 3e and **3f, S1a-S1c**, Supporting Information).

### 2.4 Validation of the inversion pipeline

The reconstruction of VEP macrostructures in sCT scans disclosed paired sagittal and lateral ‘interlocking’ protrusions on the surface of adjacent caudal and cranial VEPs (Figure 4a). These VEP protrusions (red) are short and straight, and positioned anteriorly and posteriorly along the sagittal plane, whilst those positioned laterally (blue) are elongated to form a ∼25° and ∼35° angle to the sagittal plane for the cranial and caudal VEP, respectively. The sagittal protrusions are more pronounced, and all four are taller in the cranial than the caudal VEP. Under uniaxial compression, the protrusion macrostructures on both the posterior caudal and anterior cranial VEP regions experienced concentrated first/third principal strains. The most deformed caudal and cranial regions were opposing in the anterior-posterior direction (Figure 4d and **4g**).

**Figure 4.**
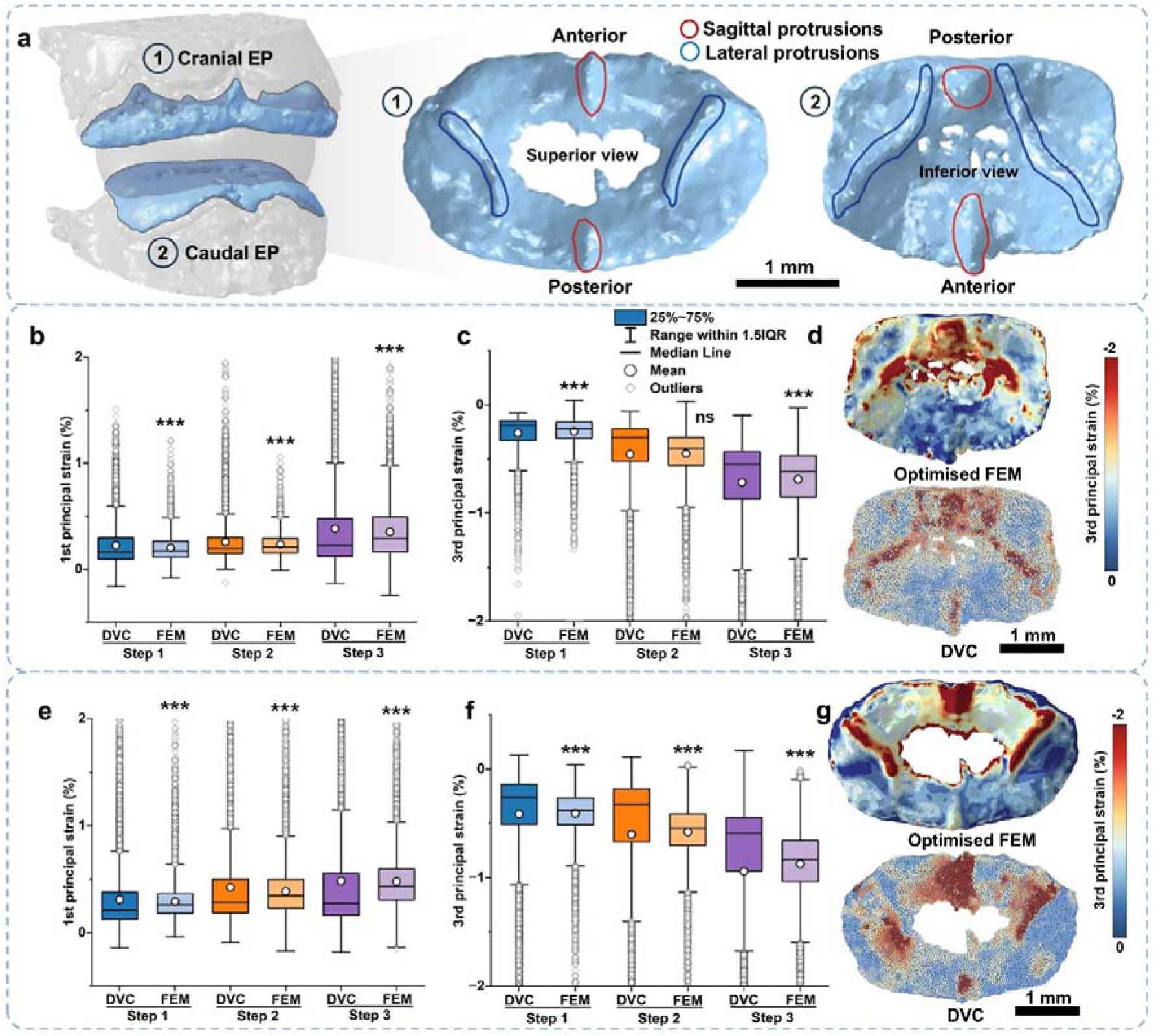
Macrostructures and strain of the caudal and cranial VEPs. **(a)** Sagittal and lateral protrusions were identified on cranial and caudal VEPs. Red circles indicate sagittal protrusions, whilst blue circles indicate lateral protrusions. The remaining flat region is defined as the ‘VEP body’; Box plots of first principal strain **(b)** and third principal strain **(c)** of caudal VEP optimised FEM and DVC point cloud reference at different loading steps; **(d)** Visualisation (inferior view) of third principal strain on caudal VEP optimised FEM and DVC point cloud reference at the second loading step (under 80 µm remote compressive displacement); Box plots of first principal strain **(e)** and third principal strain **(f)** of cranial VEP optimised FEM and DVC point cloud reference at different loading steps; **(g)** Visualisation (superior view) of third principal strain on cranial VEP optimised FEM and DVC point cloud reference at the second loading step (under 80 µm remote compressive displacement). ns: P□>□0.05; *: P□<□0.05; **: P□<□0.01; ***: P□<□0.001; Compared to DVC at the same loading steps.

Building on these structural insights, validation of pipeline efficacy shows that although the model was optimised using only the first principal strain at the second loading step (80 µm compression), the caudal VEP also generated accurate predictions at 40 µm and 120 µm compression. Compared to DVC, the distributions of FEM first principal strains showed statistical significance (p < 0.001) across three loading steps but with small η^2^ (0.2%–0.6%) (Figure 4b). The percentage difference between the mean first principal strain values of FEM and DVC were calculated, they were all smaller than the targeted 10% (e.g. 8.3% percentage difference at step 3) (**Table S1**, Supporting Information). Likewise, compared to DVC, the distributions of FEM third principal strains showed statistical significance (p < 0.001) but with small η^2^ (0.1%–0.2%) at loading step 1 and 3. No statistical significance was found at loading step 2 (p > 0.05, η^2^ = 0) (Figure 4c). The maximum difference between the mean third principal strain values across all three load steps is 5.2% at step 1 (**Table S1**, Supporting Information). This correlation in strain pattern and macrostructure aligned with visual inspection of FEM and DVC data from the caudal VEP (Figure 4d**, S1a** and **S1b**, Supporting Information).

FEM performance in the cranial VEP was similar. Compared to DVC, the distributions of FEM first and third principal strains showed statistically significant difference (p < 0.001) but with small η^2^ (0.1%–0.4%) across three loading steps (Figure 4e and **4f**). Additionally, both first and third principal strains displayed differences in mean values that were smaller than the targeted 10% at each loading step, e.g. maximum percentage difference in mean first and third principal strains between DVC and FEM were 7.9% and 7.2%, respectively, at step 2 or 3 (**Table S1**, Supporting Information). A Visual inspection revealed that cranial VEP FEM predictions exhibited a similar strain pattern to DVC (Figure 4g and **Figure S2**, Supporting Information).

FEM in both caudal and cranial VEP were, however, partially inconsistent with DVC in the high-strain regions, with 2–4% of the predictions significantly underestimating first and third principal strains, but with a strong correlation still retained at all loading steps (>95% of nodes with 95% limit of agreement **Table 1**).

**Table 1.** Statistics of the Bland□Altman plots at different compression steps.

| Samples | Type of strain | Compression steps | Nodes within 95% limits of agreement (%) | Overestimated nodes (%) | Underestimated nodes (%) |
| --- | --- | --- | --- | --- | --- |
| Caudal VEP | First principal strain | 1 | 96.1 | 0.5 | 3.4 |
|  |  | 2 | 96.5 | 1.1 | 2.4 |
|  |  | 3 | 95.4 | 0.9 | 3.7 |
|  | Third principal strain | 1 | 95.7 | 1.3 | 3.0 |
|  |  | 2 | 96.0 | 1.0 | 3.0 |
|  |  | 3 | 95.5 | 0.5 | 4.0 |
| Cranial VEP | First principal strain | 1 | 95.3 | 1.0 | 3.7 |
|  |  | 2 | 95.7 | 0.8 | 3.5 |
|  |  | 3 | 95.8 | 0.7 | 3.5 |
|  | Third principal strain | 1 | 95.7 | 0.3 | 4.0 |
|  |  | 2 | 96.0 | 0.6 | 3.4 |
|  |  | 3 | 96.2 | 0.7 | 3.1 |

Note: Step 1: 40 $\mu\text{m}$ remote compressive strain; Step 2: 80 $\mu\text{m}$ remote compressive strain; Step 3: 120 $\mu\text{m}$ remote compressive strain.

### 2.5 Consistency of the pipeline across different specimens

The pipeline consistency was tested on three different rat lumbar FEMs. Figure 5a and **5b** illustrate the obtained sCT imaging data of three rat specimens and the FE models constructed from sCT, where we found variations in VEP thickness, trabecula structure, and overall morphology. Differences in sample preparation angles and dissection orientations were also observed. Although there are some objective or unavoidable inconsistencies between the specimens, the pipeline successfully converged on each case. The optimised VEP material properties for the three specimens vary, and have a mean elastic modulus of 144 ± 18 MPa and Poisson’s ratio of 0.28 ± 0.04 (Figure 5d, **Table 2**). There are 186 ± 44 local minima identified, and the RMSE values for the global minima of each case remain low (2.33 ± 0.17 × 10□³). On the optimised FEM, strain distribution analysis (Figure 5c) under 80 µm remote compressive displacement shows that the third principal strain in different VEP regions across the three FEMs follows similar trends. Specifically, in cranial VEPs of Lumbar 1, 2, and 3, higher strain values with statistical significance (p < 0.01 or < 0.001, η^2^ = 17.5%, 37.8%, and 41.8%, respectively) were observed in both lateral and sagittal protrusions, compared to the VEP body. In caudal VEPs of Lumbar 1, 2, and 3, similar higher strain values with statistical significance (p < 0.05 or < 0.001, η^2^ = 9.7%, 6.7%, and 6.2%, respectively) were also observed in both lateral and sagittal protrusions, compared to the VEP body.

**Figure 5.**
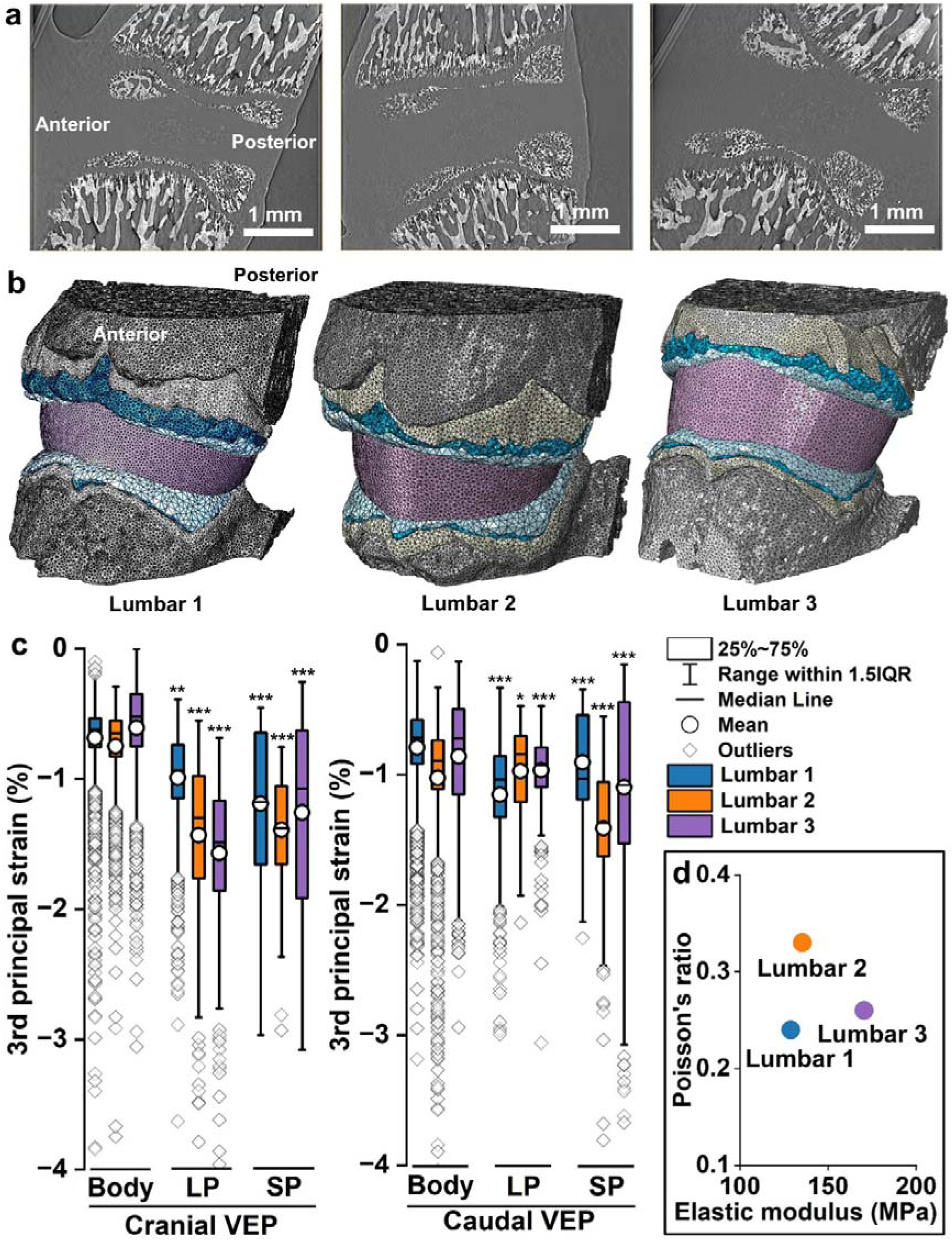
Comparison of FEMs and third principal strain distribution across three different rat lumbar segments. **(a)** sCT imaging of three rat lumbar segments used for model construction; **(b)** Three FEMs as inputs of the pipeline for determining their VEPs’ elastic modulus and Poisson’s ratio; **(c)** Regional third principal strain distribution analysis on the three optimised FEMs; **(d)** Optimised combinations of elastic modulus and Poisson’s ratio for three VEPs. ns: P□>□0.05; *: P□<□0.05; **: P□<□0.01; ***: P□<□0.001; Compared to the VEP body in the same lumbar.

**Table 2.** Optimisation results of elastic modulus and Poisson’s ratio across three rat VEPs.

| Rat | Elastic modulus (MPa) | Poisson's ratio | RMSE of Global minima ($\times 10^{-3}$) | Total number of local minima |
| --- | --- | --- | --- | --- |
| 1 | 129 | 0.24 | 2.09 | 129 |
| 2 | 135 | 0.33 | 2.50 | 194 |
| 3 | 170 | 0.26 | 2.40 | 235 |

## 3 Discussion

This study was focused on rat VEPs that were used to develop and validate an FEM inversion pipeline. The data indicates that the pipeline enables FEM predictions to achieve a high level of agreement with DVC experimental measurements. Despite sample variations, the pipeline reliably converged on a narrow range of material properties, with an elastic modulus of 145 ± 18 MPa and a Poisson’s ratio of 0.28 ± 0.04.

The combination of high-resolution sCT, DVC and *in situ* mechanical testing has exhibited novelty in quantifying 3D full-field strains with nanoscale precision[22, 33]. For spine/whole joints, this approach has been applied to mouse whole knee joints[22], rat whole lumbar intervertebral discs[20–21, 23, 34], and human thoracic vertebrae cylinders[35]. While sCT imaging provides high spatial resolution for structural details and DVC is invaluable for comparing local and global strains, most studies have focused only on relatively small, homogeneous samples. Application of sCT-based *in situ* DVC in intact joint biomechanics, including both soft and hard tissues, is still at an early stage. To our knowledge, rat spine studies utilising sCT-based *in situ* DVC have only been reported by Parmenter *et al*.[23, 36] (preprint/conference presentation) and Disney *et al*.[20–21, 34]. The latter elegantly correlated structural responses in collagen bundles to uniaxial compression, and reported differences in annulus fibrosus compliance of the different intervertebral disc regions[21],. The former showed that local strain distribution varies regionally at the annulus fibrosus-endplate junction, and elucidated the relationships between microstructure and mechanics, nanoscale mineral structure, and molecular level prestrain in VEP.

Despite the advantages of sCT-based *in situ* DVC, bidding processes and long waiting times deter access to sCT facilities[37]. Bespoke sample holders for *in situ* mechanical load application also need to be designed and manufactured and, with cost and size of a multiaxial mechanical tester, it is not surprising that our experiments were all performed on simpler uniaxial compression/tension devices. sCT-based *in situ* DVC is also limited by animal model availability. It is generally acknowledged in bone research that larger sheep and dogs, more closely resemble human biomechanics and pathology[38]. These are often more expensive requiring complex housing facilities and management, which limits their use. For these reasons, FEM has promise in providing new insights into VEP biomechanics, as it not only allows elucidation of global VEP strain/stress but can also simulate the influence of different motions and disease conditions without much additional investment. It is likewise important to appreciate that a set of DVC data is indispensable for constructing and validating the FEM before the model can be used for further simulations. Previous studies have used DVC to validate FEM of whole bones[39–42], however these models rely on known material properties of the modelled system, which are not always available.

Material properties significantly affect FEM prediction accuracy, but accurate material properties for complex multi-tissue systems are often unavailable. Thus, one of our aims was to develop a pipeline that allows global material properties of a complex rat VEP, consisting of both bone and cartilage, to be adjusted in line with DVC data. There is a lack of existing data for material properties of the spine in animal models, so human material properties from literature were used in our rat lumbar FEM as a starting assumption. Existing data for apparent moduli from mouse[43] and rat[43–44] spine segments are indeed in the same order of magnitude as reported moduli of human spines[43] and, while some studies have reported similar inverse methods on homogeneous materials[29, 45], our study is the first to apply this method to a functionally intact joint. More details regarding the rationality can be found in **Supplementary Methods** and **Supplementary Note 1.** Our pipeline also affords an advancement in its use of a full-field node-to-node spatially registered reference and its scope to vary material properties.

Prior to performing the pipeline, our mesh convergence test confirmed the reasonable mesh size for stable predictions in each FEM part. The sensitivity test suggests that the mechanical response of VEPs is affected by both direct and indirect contact structures, with the greatest deviation (18%) in VEP strain caused by annulus fibrosus. Such variability could introduce uncertainty into the FEM predictions, potentially leading to discrepancies in the optimised elastic modulus and Poisson’s ratio. Additionally, we estimated the impact of utilising elastic or viscoelastic material properties for intervertebral discs under our quasi-static conditions (See **Supplementary Methods** and **Supplementary Note 2** for details). It is important to note that our pipeline is not aimed at providing ground truth material properties but an equivalent estimation, ensuring that the predicted mechanical response aligns with experimentally observed strain distributions. Therefore, our pipeline can dynamically adjust the estimation within a reasonable predefined range. The spatial registration effectively preserved the original DVC point cloud strain features with minimal deviations. While the Welch’s ANOVA revealed that smoothing extreme values led to significant differences in the distribution of principal strains (p < 0.001), the effect size was small (η² = 0.5%). This indicated that although differences were statistically detectable due to the large sample size, they were practically negligible. This was further evidenced by minor peak shift (up to 0.03% absolute strain value, within the overall accuracy of DVC measurements) and low Kullback-Leibler divergence values (up to 0.89).

Using the registered DVC and FEM, our pipeline integrates varied 3D strain values in different VEP regions, while identifying global elastic modulus and Poisson’s ratio in RMSE from local optima. Interestingly, the material properties used in prior FEM studies (**Table S6**, Supporting Information) of humans exposed marked inconsistencies in properties both within and across spine segments. Comparing the outcomes of our pipeline with reported human VEP material properties, we find that global optimal values for both rat VEP equivalent elastic modulus and Poisson’s ratio falls within the human VEP range (144.9 ± 18.2 MPa in rats vs. 24-6,000 MPa in humans, and 0.28 ± 0.04 in rats vs. 0.25-0.4 in humans). It is vital, however, to highlight that determination of these parameters is complicated by the inhomogeneous, anisotropic, viscoelastic, and nonlinear[46] properties of biological tissues. Determination of VEP material properties can be further complicated by sample selection (spine segments, anatomical differences, donor age) and preparation (minor alignment deviations), measurement standards (macroscale properties[47], microscale indentation[48], or bone mineral density-dependent CT grayscale prediction[48]) and region measured[49]. Nevertheless, the pipeline demonstrated consistent VEP strain distribution trends across all models with median to large effect size (η² = 6.2%–41.8%), indicating that the observed variability primarily reflects biological differences rather than methodological inconsistencies. Additionally, note that the due to the element size (minimum edge length 32.5 μm) used for VEP, all elements incorporate both bony VEP and unmineralised components (**Figure S8**, Supporting Information), providing a homogenised elastic modulus value. This likely contributes to the relatively lower apparent modulus and Poisson’s ratio obtained from our rat VEP model, compared to some experimental measurements in human and rabbit samples using indentation. Because the optimised values are homogenised, they are still appropriate for finer meshes as long as spatially varying properties are not applied. They would not be appropriate if spatially resolved moduli were available and the aim was to model their distribution explicitly at sub-element scale. However, the values are appropriate for coarser meshes. Furthermore, higher apparent values measured by universal testing machines may be attributed to three additional factors: (i) In mechanical testing, real specimens may become compacted under load, leading to overestimated apparent moduli; (ii) Species-specific differences; (iii) Age-related effects, since human specimens are usually obtained from older donors with mature endplate calcification, whereas our specimens from 8-week-old rats (roughly equivalent to 20-year-old humans) were less calcified. The sensitivity test also indicates that the modulus/Poisson’s ratio used for other structures will affect the predicted VEP material properties. Such factors can clearly contribute to significant variation between studies. Importantly, our optimal VEP material properties resulted in good statistical and spatial consistency of strains with DVC data to validate the robustness of our FEM in the unsupervised strain prediction in this complex load transfer system. Although the statistical analysis reveals significant differences (p < 0.001) between FEM and DVC strain distributions at node level, the small effect sizes (η² = 0.1%–0.6%) indicate that these differences have minimal practical significance. This interpretation is supported by the <10% differences between mean strain values and a >95% agreement using the Bland-Altman analysis across all loading steps at whole model level.

We noted that FEM predictions had 2–4% strain values above/below the 95% limits of agreement with DVC data. This may reflect differences between FEM and DVC algorithms, as DVC tracks the natural microstructure-related grayscale variations in microstructural features [50] and can reflect effects such as trabecular failure. Previous studies have reported a load-induced crack development and failure of fine trabeculae (∼100 μm) leading to significantly raised DVC strain values[51]. Our FEM was informed by binned sCT images that averaged fine trabeculae which, in contrast, did not consider damage, and thus will not be able to predict failure, and whilst failure has been reported to occur with similar ∼1% compressive strain in bovine and human trabeculae[52], nonlinear and failure response was not included in our FEM due to a lack of data in the rat. These factors may explain the increasing discrepancy observed in our dataset, which gradually becomes more significant at higher strains approaching and exceeding 1% (Figure 3d). We underline that our FEM does not capture all biomechanical behaviours but instead estimates physiology closely using only moderate computational power. The inversion pipeline was indeed effective in a complicated system composed of ‘interacting’ parts with divergent material properties and can use material properties from humans. DVC-based FEM material property inversion could capture the range of strain values and their spatial distribution, which is critical for more detailed and robust analyses in different species and individual parts of the spine. Our pipeline can also be adapted to conventional lab-based microCT datasets. However, its application may be limited to hard tissues, as the soft tissues have poor contrast in conventional microCT. Beyond this methodology development study using small animals, clinical MRI-based DVC has been used on human intervertebral discs to show strain localisation mechanisms[53], demonstrating the promise of transferring our inversion pipeline to large animals and humans.

There are further limitations in this study. Our coarse mesh caused inevitable merging of some fine trabeculae (∼10 μm) into larger structures (∼30 μm) and thus less concentrated strains. In addition, potential sources of error may arise from semi-subjective segmentation inaccuracies, particularly at complex interfaces such as the VEP boundaries. A highly detailed model should be created and run on a computer cluster for comparison in the future. Although uniaxial loading was carefully controlled and recorded, slight misalignments or actuator compliance may have introduced minor deviations in the true boundary conditions. The FEM included only the isotropic elastic modulus and Poisson’s ratio, while plasticity and damage under load were not considered due to a lack of experimental data. The intervertebral disc part of the FEM is more simplified than other parts. Although a lamellar structure and orientation-dependent properties were considered, the ± 45° fibre orientation was an approximation and they did not completely resolve the individual annulus fibrosus collagen bundles and their varied orientation within the same lamella[21], which limited us to further studying interaction at annulus fibrosus-endplate junctions and the load transfer path from one VEP *via* the collagen bundles to another. It should also be noted that VEP structure differs among species[54] and that rodents and humans have distinct forms of locomotion. The rat model studied herein does not have direct translation to humans[55]. Hence, more consideration should be given when animal models are used to study human diseases.

## 4 Conclusions

In summary, our study established an inversion pipeline for optimising the material properties of rat lumbar VEPs in a spine motion segment FEM. The optimised FEM displayed strain values and distributions which are in good agreement with the DVC measurements. VEP macrostructures linked to strain concentrations were identified on both cranial and caudal VEPs. Although we demonstrated these methodologies on small animals, they have the potential to be scalable to large animals and humans, and we are planning experiments for further validation.

## 5 Experimental Section

### Description of Methodology

We developed an FEM/DVC pipeline for optimised determination of homogenised values of elastic modulus (E) and Poisson’s ratio (ν) in the VEP using DVC data (see **Figure S9,** Supporting Information). Briefly, three separate initial inputs were provided to the optimisation pipeline: 1) an image-based rat lumbar FEM mesh with high resolution microstructures obtained from static sCT scans; 2) DVC strains obtained from *in situ* quasi-dynamic sCT scans; and 3) default VEP isotropic elastic modulus and Poisson’s ratio values from literature as a starting point of optimisation. The optimisation pipeline determined the optimal combination of elastic modulus and Poisson’s ratio in the VEP, which is an output giving the smallest RMSE between FEM predictions and DVC strains. The caudal DVC dataset at displacement of 80 µm was used for the pipeline, while the cranial DVC datasets at displacement of 40 µm, 80 µm, and 120 µm and the caudal datasets at displacement of 40 µm and 120 µm were reserved as independent validation datasets. The pipeline was then applied to the other two rat spine specimens to validate the optimised values of elastic modulus and Poisson’s ratio.

### Experimental animals, *in situ* uniaxial compression, sCT, image reconstruction and DVC

The ethical approval for the use of experimental animals and tissue collection procedures was approved by the University of Manchester Animal Welfare and Ethics Review Board and followed the University’s Animal Research Policy and the UK Animal (Scientific Procedures) Act of 1986 under the Home Office Licence (#70/8858 or I045CA465). The two male, 8-week-old Sprague Dawley rats used for this study were housed at the University of Manchester Biomedical Sciences Facility. Following euthanasia, three lumbar segments were dissected from the two rats, with one contributing a single lumbar vertebral pair (L4/5, Lumbar 1) for the establishment of the pipeline, while the other contributed two vertebral pairs (L2/3, Lumbar 2 and L3/4, Lumbar 3) for pipeline validation. These three lumbar vertebral pairs were chosen because of sample quality and complete force and imaging data across all loading steps. *In situ* sCT, image reconstruction, and DVC of rat lumbar segments were performed as described in [36]. Briefly, sCT imaging was performed at the Diamond Light Source beamline I13-2 [50] using a pink beam (mean energy 27 keV), voxel size of 1.625 µm, and 1800 projections per scan. The rat spine segment, consisting of an intervertebral disc with half-vertebrae either side and posterior elements removed, was placed in a loading rig (Figure 1a) with a 1N pre-load applied (equivalent to ∼40 % of the animal’s body weight), followed by three displacement steps of 40 µm at a constant remote displacement rate of 0.1 mm/min. Four sCT scans were taken per sample, 1200 s after the application of each loading step to allow for stress relaxation. Each scan took 3 min and consisted of 1800 projections acquired using fly scanning, with an exposure time of 0.1 s/projection. The total duration of each in situ experiment was ∼100 min, from sample mounting in the loading rig to the acquisition of the final scan. sCT scan reconstruction was conducted using Savu[56].

DVC analysis was performed using source code supported by CCPi (Collaborative Computational Project in Tomographic Imaging, https://tomographicimaging.github.io/iDVC/, version 21.1.0), using a local approach with flexible point cloud specification. Microstructure specific point clouds of the VEPs were generated from meshes of the mineralised tissue structure, with distance between points set 4.5 voxels (7.3 µm), and DVC was performed using a sub-volume diameter of 30 voxels (49 µm), 2000 sampling points per volume, and 12 degrees of freedom. To enable strain measurements at a comparable resolution to the FEM, the displacement field point cloud was randomly down sampled to 2.5 % the number of points, reducing the number of points from ∼1,000,000 to ∼25,000, and increasing the mean minimum distance between points from 4.5 voxels (7.3 µm) to 14.6 voxels (23.7 µm). Strains were calculated by polynomial fitting to the 25 closest points, resulting in a mean strain window radius of 42.7 voxels (69.4 µm), and a virtual strain gauge of 115.4 voxels (187.5 µm). Accuracy of the measured strains was determined from a standard DVC zero-strain analysis, where two scans were taken of the same sample without applying any strain. This gave a measured strain accuracy of 0.031 % and precision of 0.034 % (calculated from the mean absolute error and standard deviation of error respectively).

### FE model of the rat lumbar segment

The FE model geometry of the rat lumbar vertebral segment was defined from the acquired sCT images. The sCT field of view captured the entire intervertebral disc, VEPs, and approximately 1 mm of the vertebrae either side of the intervertebral disc. The reconstructed images were first binned by four using Avizo 3D (v. 2023.2, Thermo Fisher Scientific, Waltham, US), increasing the voxel size from 1.6 µm to 6.5 µm. After that, the segmentation module in Avizo 3D was used for segmenting the volumes of the vertebrae, growth plates, CEPs, and VEPs. Segmentation was based on the anatomical boundary of the structures. A mesh wrapping only the VEPs was defined as the region for DVC and FEM process. Before modelling, an image binning strategy was applied to mineralised tissues in the VEPs and vertebrae, which merged very small trabeculae (∼10 μm), but preserved larger trabeculae (∼50 μm), to necessitate a trade-off between model resolution and computational time.

For the annulus fibrosus and nucleus pulposus, 3D models of five annulus fibrosus lamellae and their associated nucleus pulposi were created according to their boundary and shape in the sCT images using Cinema 4D (v. 2023, Maxon, Bad Homburg, Germany). All of the parts were then re-imported into Avizo 3D and converted into surface data with a constrained minimum edge length according to the convergence study. After that, all of the surface data were converted into tetrahedral elements. A model comprising vertebrae, growth plates, CEPs, VEPs, five annulus fibrosus layers, and nucleus pulposus (Figure 1c-1g) was constructed. The tetrahedral elements were then exported as INP files adapted to Abaqus CAE (v. 2023, Simulia, Providence, US), which was used to perform finite element analysis on the strain and stress on the spine. After the materials were imported into Abaqus/Standard, the material properties were determined, as detailed in **Table 3** (Supporting Information) and **Table 4** (Supporting Information). Local coordinate systems were defined in annulus fibrosus lamellae to mimic anisotropy in each lamella (**Table 4**, Supporting Information). Our previous work showed that collagen bundles in annulus fibrosus lamellae have alternating ± 30° or ± 60° orientations, depending on their location in the intervertebral disc[20–21]. To represent the anisotropic orientation in each lamella, an average alternating ± 45° material orientations were applied to the annulus fibrosus in the FEM. The parts were assembled into an integrated model according to their original spatial relationship, the corresponding interactions were set to connect the parts, and surface-to-surface contact was applied between the annulus fibrosus lamellae and the innermost annulus fibrosus lamella and NP (**Figure S10**, Supporting Information). The highest and the lowest planes of the lumbar segment FEM were kinematically coupled with two reference points (RP1 and RP2, corresponding to top and bottom surfaces in Figure 1c respectively) which were located at the centre of the planes. All boundary conditions were applied to these two reference points which have constrained all degrees of freedom to the planes. The boundary conditions were the same as those in the experiment. Briefly, RP1 was encastred to replicate the experiment, where the top of the spine segment was glued to the compression stage, while the lowest plane of the caudal vertebra was subjected to compressive displacement *via* three 40 µm steps. Displacement was applied at RP2 along the Z direction, with constrained zero displacement in X and Y directions, and no rotations about any of the axes. The strain and stress over the FEM were then calculated in the implicit solver. The FEMs of the other two rat spine segments for validating the pipeline consistency followed the same methods above.

**Table 3.**
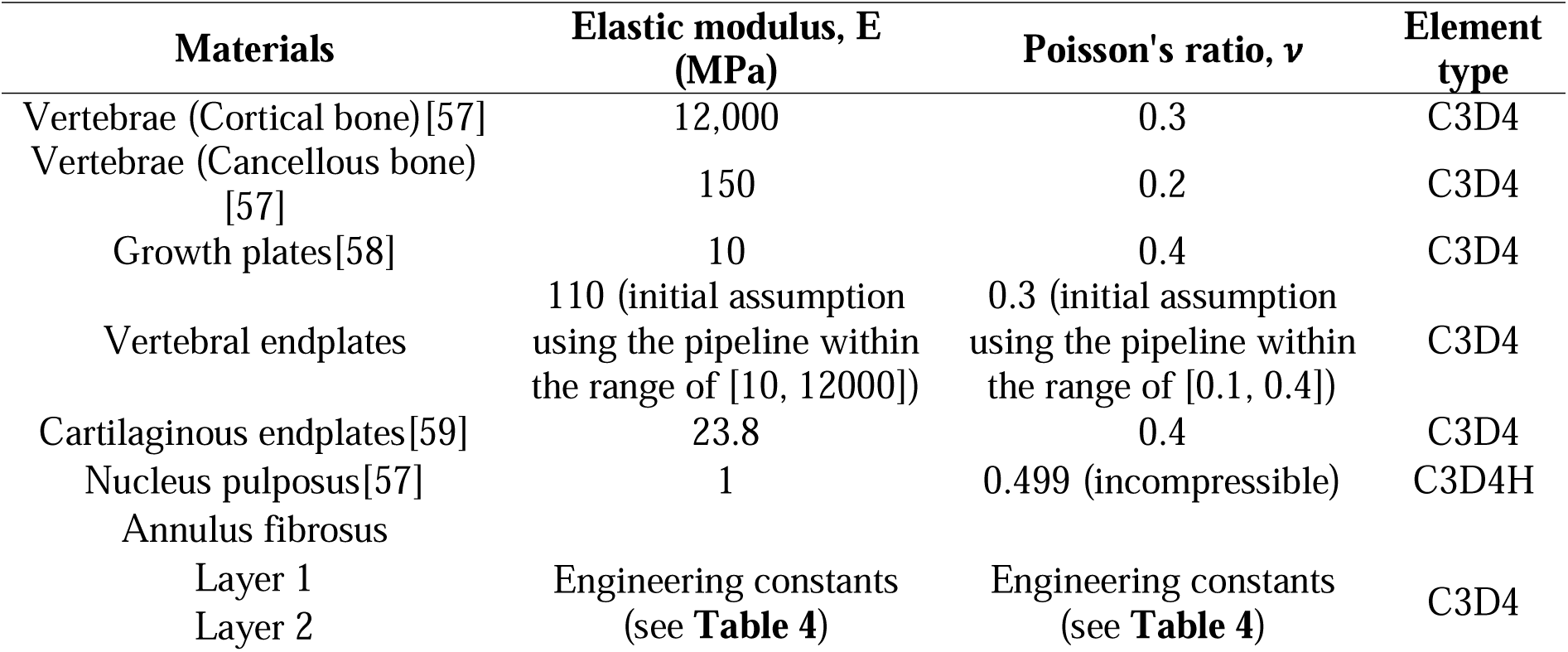
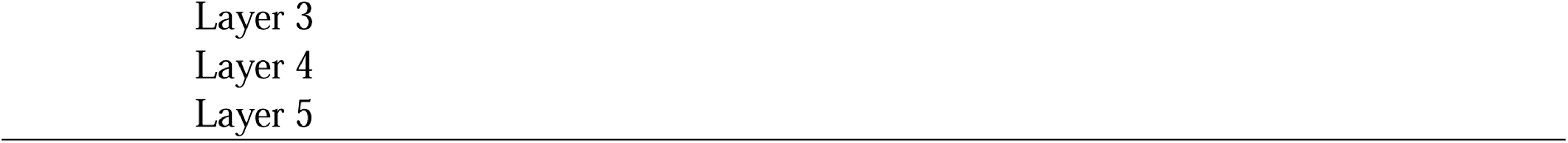
Material properties of the finite element model of rat lumbar spine segments according to the references with minor modifications.

**Table 4.**
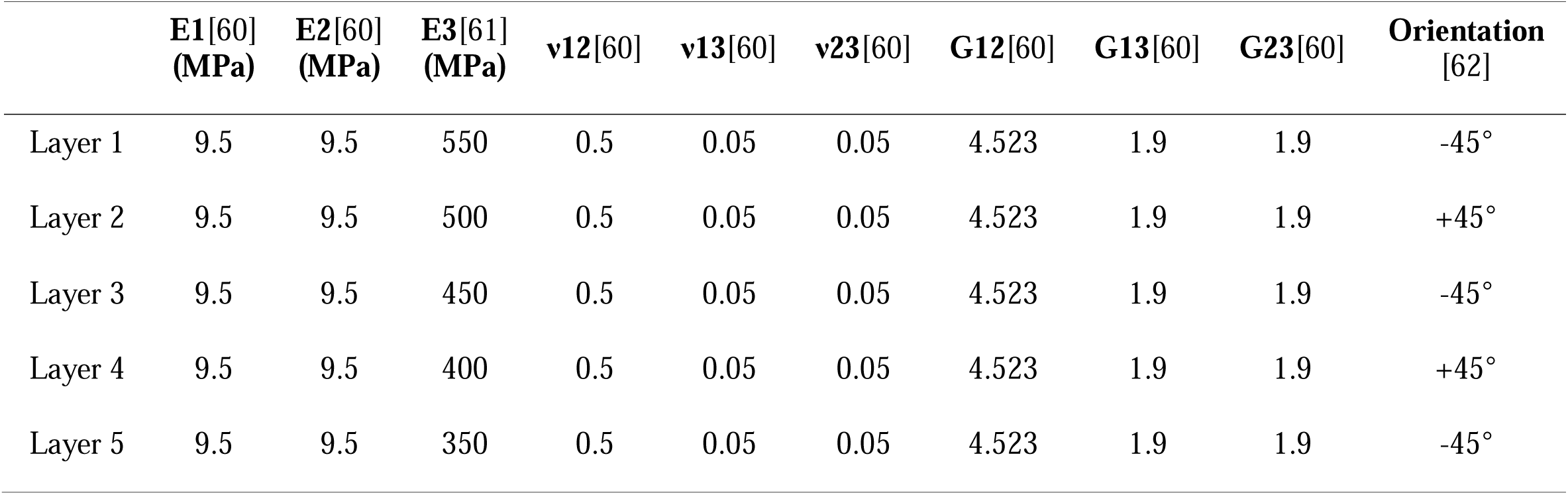
Anisotropic engineering constants of annulus fibrosus lamellae of the finite element model.

### Mesh convergence study of FEM

A mesh convergence study was conducted on all FEM parts. Only one part was subjected to the convergence test each time. A series of tetrahedral elements with different minimum edge lengths (24.4 μm, 32.5 μm, 40.6 μm, 48.8 μm, 56.9 μm, 65 μm, 73.1 μm, and 81.3 μm) were created in Avizo 3D following the description above. The edge length of 40.6 μm was selected as the default for all parts, and the edge length of the adjusted part was sequentially varied in the order listed above. They were then exported as INP files and imported into Abaqus CAE to replace the original VEPs FEM. All elements were linear tetrahedral with four nodes and one integration point. Accordingly, combination material properties (E = 110 MPa, ν = 0.3) were initially assigned to the VEP by the MATLAB Global Optimisation Toolbox with boundary condition of 80 μm compressive displacement, and static analysis conducted with the implicit solver. After obtaining the strain data on the VEP FEM, the average first principal strain at the nodes was calculated. Plots of the average first principal strain vs the number of elements on both VEPs were generated using Origin (v. 2023b, Originlab, Northampton, US). When a threshold value of 1% of the rate of change of the average first principal strain with element size is achieved, the mesh was considered converged.

### Sensitivity test of FEM

A sensitivity test was conducted by adjusting the elastic modulus of all FEM parts except the VEPs (E = 110 MPa, ν = 0.3). Each FEM part was tested individually while keeping the material properties of other parts constant. The values were selected based on the maximum and minimum reported in the literature (**Table S7**, Supporting Information) to estimate the potential range of influence on both the adjusted part itself and the VEP strain distribution. Since the Poisson’s ratio of biological tissues is generally assumed rather than directly measured, only the elastic modulus was varied in this test. The simulations were performed using the same boundary conditions as the convergence test with an applied 80 μm compressive displacement. For each case, the first principal strain averaged from the adjusted parts and the VEP were assessed.

### Registration of the DVC point cloud to FEM nodes

Prior to the global optimisation pipeline, DVC point clouds were registered to FEM nodes *via* a bespoke code in MATLAB (v. R2023b, MathWorks, Natick, US). The original caudal and cranial DVC meshes contained 23,529 and 29,187 nodes, respectively. First, the original DVC mesh (Mesh 1) was replaced by a new DVC mesh (Mesh 2) with the same node number and node spatial position referenced to the FEM mesh (Mesh 3). On Mesh 2, the nodes were then assigned values of first, second, and third principal strain from Mesh 1 following two principles: 1) if the Mesh 2 nodes were within the Mesh 1 domains, natural neighbour interpolation – a method based on Voronoi tessellation and suitable for irregularly spaced data – was used to generate strain values for Mesh 2 nodes; 2) if the Mesh 2 nodes were outside the Mesh 1 domains and natural neighbour interpolation could not be applied, extrapolation by the nearest Mesh 1 nodes was performed to avoid potential error accumulation from simple linear extrapolation (**Figure S11**, Supporting Information). To validate the comparability, Mesh 2 was then compared with Mesh 1 on statistical and spatial distributions of first, second, and third principal strain values at both sub-region and single point levels.

### Sub-region-based spatial registration error distribution analysis

To evaluate the consistency of DVC strain fields before and after spatial registration, we firstly performed a sub-region-based error distribution analysis. The strain data were processed by splitting the spatial domain into sub-regions of cubes with 32.5 µm edge length, ensuring a consistent spatial resolution for comparison. Within each sub-region, the first, second, and third principal strains of all points were averaged to obtain representative strain values. The absolute error in principal strains was then calculated for each sub-region by comparing the registered and unregistered DVC strain fields. The relative frequency of these errors was visualised using histogram.

### Single point-based Kullback–Leibler divergence and Kernel density estimation

Kullback–Leibler divergence was performed to evaluate the variation when mapping first, second, and third principal strain values from Mesh 1 to Mesh 2. The divergence value ranges from zero to positive infinity. Generally, two identical datasets give a divergence value of zero. In terms of two varied datasets, a smaller divergence value indicates greater similarity between the two distributions. This approach provides a quantitative measure of the interpolation and extrapolation accuracy. Prior to calculating the Kullback–Leibler divergence, Kernel density estimation was applied to approximate the probability density functions of first, second, and third principal strain distributions for both meshes. Kernel density estimation was used to ensure smooth and continuous distributions, with the same evaluation points used for both meshes to allow direct comparison. The Kernel density functions were normalised to represent valid probability distributions, and an epsilon value □ = 1 × 10^-10^ was specifically added to data points where the density was zero to avoid log(0) or divided by zero error. The Kullback–Leibler divergence was calculated using **Equation 1**:

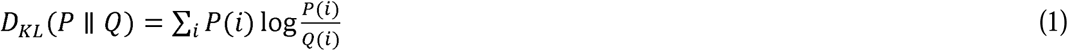

where

is the probability densities of Mesh 2.
is the probability densities of Mesh 1.

### Global optimisation pipeline

The global optimisation pipeline was established on integrated bespoke codes on the platforms of MATLAB and Abaqus CAE built-in Python. The first principal strain on the caudal VEP was utilised for the pipeline. **Figure S12** (Supporting Information) outlines the workflow. Briefly, a bespoke MATLAB function was used to modify the elastic modulus and Poisson’s ratio of the caudal VEP FEM in the INP file of Abaqus CAE. The adjustment ranges of the elastic modulus and Poisson’s ratio were set to [10, 12,000] MPa and [0.1, 0.4], respectively. This range is based on a reasonable assumption that the VEPs are not stiffer than the cortical bone of vertebrae nor softer than the growth plates. Then, the Abaqus Standard was launched, and the INP file containing all the information on the parts, models, interactions, and boundary conditions (80 µm compressive displacement) was imported into Abaqus. After that, the models were solved using the Abaqus built-in implicit solver. The first principal strain data of the caudal VEP FEM were extracted by another bespoke MATLAB function for comparison with the first principal strain data of DVC in node-to-node form. Deviations from each pair of FEM-DVC nodes contribute to the final RMSE (**Equation 2**), which reflects the degree of difference between two paired datasets. The MATLAB Global Optimisation Toolbox using gradient descent-based algorithms was used to analyse the RMSE values and determine whether they had achieved the global minimum. If not, the combination of the elastic modulus and Poisson’s ratio would be automatically adjusted and rerun above the workflow; otherwise, the global optimisation pipeline would be terminated, and the global optimal combination of the elastic modulus and Poisson’s ratio and corresponding RMSE would be printed. The visualisation of the optimisation results in the full range was plotted using Origin.

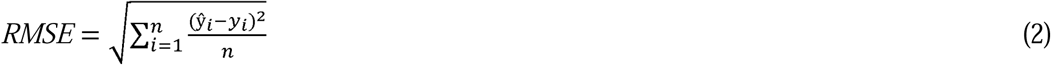

where

*ŷ* is the FEM-predicted value.
*y* is the DVC measured value.
*n* is the number of node pairs.

### Validation of the optimisation pipeline

In the caudal VEP, once FEM-predicted first principal strain was matched to DVC measurements, the optimisation pipeline efficacy was further validated by longitudinally comparing FEM/DVC first and third principal strains on caudal/cranial VEP from the preload (1N) to three loading steps (40 µm, 80 µm and 120 µm strain of full intervertebral disc height). A strain difference of less than 10%[63] between FEM and DVC was considered as a good match.

### Linear regression analysis of paired points

Linear regression was performed to analyse the agreement of the average first principal strains between the FEM predictions and DVC measurements under 80 µm remote compressive displacement on the caudal VEP. All paired points were extracted from FEM and DVC references, and the agreement was assessed utilising linear regression in SPSS Statistics (v.29.0.0.0, IBM, New York, US) and then plotted utilising Origin.

### Bland–Altman plot

Data normality was tested and Bland–Altman plots were constructed to study the agreement of first principal strain under 80 μm compressive displacement on the caudal VEP between the FEM predictions and DVC measurements. The Bland–Altman plots were created based on the averages (x-axis, **Equation 3**) and differences (y-axis, **Equation 4**) of first principal strains between the paired FEM and the DVC data, followed by computing the mean and standard deviation of these differences to establish 95% limits of agreement, defined as the mean difference plus or minus 1.96 times the SD[31–32]. In the situation where two independent measurement methods have good agreement, over 95% of the data points should fall within the range of limits of agreement. The visualisation of the Bland–Altman plots was performed using Origin.

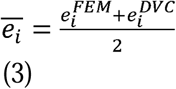

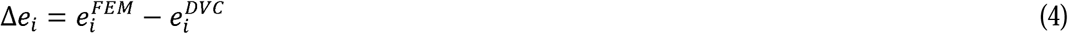

where

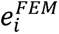 is the FEM-predicted first principal strain.
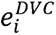 is the DVC measured first principal strain.

### Statistical analysis

All statistical analyses were performed using SPSS Statistics and visualised using Origin. Levene’s test was used to assess homogeneity of variances. When variances were equal, standard one-way ANOVA was applied; otherwise, Welch’s ANOVA was used as a robust alternative. Effect sizes (η²) were calculated to assess the practical significance of group differences, with values interpreted as small (1%), medium (6%), and large (14%). To compare the distribution of principal strains before and after spatial registration, Welch’s ANOVA with η² was used. As complementary quantifications for spatial registration accuracy, sub-region-based absolute error distribution, kernel density estimation, and Kullback–Leibler divergence were employed. Differences between FEM predictions and DVC measurements were assessed using Welch’s ANOVA with η², Bland–Altman plots, and linear regression analysis (R²). Regional strain variations in the VEPs of Lumbar 1, 2, and 3 were compared using Welch’s ANOVA with η². For post hoc comparisons, LSD tests were used under the assumption of equal variances, and Tamhane’s T2 test was applied when variances were unequal. Where applicable, data are presented as mean ± standard deviation, with 95% confidence intervals. A significance level of p < 0.05 was considered statistically significant.

## Data Availability Statement

Example sCT scan data are available from the UCL Research Data Repository (DOI: 10.5522/04/26789212.v1). The FE models of the three rat lumbars, and full codes of the inversion pipeline are available on Zenodo (DOI: 10.5281/zenodo.15704567).

## Acknowledgements

We sincerely thank Prof Michael J Sherratt for providing the experimental animals for this study. This work was supported by grants EP/V011235/1, EP/V011006/1, and EP/V011383/1 from the UK-Engineering and Physical Sciences Research Council, MR/V033506/1 from the UK-Medical Research Council, grant CZIF2021-006424 from the Chan Zuckerberg Initiative Foundation, and grant 2022-316777 from the Chan Zuckerberg Initiative DAF, an advised fund of Silicon Valley Community Foundation. We gratefully acknowledge Beamtime SM29784 at Beamline I13-2, Diamond Light Source; and Beamtime MD-1290 and MD-1389 at Beamline BM18, European Synchrotron Radiation Facility. The authors gratefully acknowledge Joseph Brunet’s beneficial comments on the modelling.

## Ethics approval statement

The ethics for using the experimental animals and tissue collection procedures was approved by the University of Manchester Animal Welfare and Ethics Review Board and followed the University’s Animal Research Policy and the UK Animal (Scientific Procedures) Act of 1986 under the Home Office Licence (#70/8858 or I045CA465).

## Author Contributions

Conception and design of the study: P.D.L. and H.S.G. Acquisition of data: J.C., A.P., A.S., E.N., S.M., and A.A.P. Interpretation of data, revision of the manuscript, final approval and agreement to be accountable for all aspects of the work: all authors. Drafting of the manuscript: J.C., A.P., A.S., and P.D.L.

