## Supplementary Materials for "Synchrotron Tomography-Based Inversion Pipeline for Estimating Elastic Properties of Rat Vertebral Endplate Finite Element Models"

**Synchrotron Tomography-Based Finite Element Analysis of Vertebral Endplate Loading Reveals Functional Roles for Architectural Features**

Jishizhan Chen^1^, Alissa Parmenter^1,2^, Aikta Sharma^1^, Elis Newham^3^, Eral Bele^1^, Sebastian Marussi^1,2^, Andrew A Pitsillides^4^, Nick J Terrill^5^, Christopher Mitchell^6^, Himadri S Gupta^3,*^, Peter D Lee^1,2,*^

^1^ Mechanical Engineering, University College London, Torrington Place, London WC1E 7JE, UK

^2^ Research Complex at Harwell, Rutherford Appleton Laboratory, Oxfordshire OX11 0FA, UK

^3^ School of Engineering and Materials Sciences, Queen Mary University of London, London E1 4NS, UK

^4^ Department of Veterinary Basic Sciences, Royal Veterinary College, Royal College Street, London NW1 0TU, UK

^5^ Diamond Light Source, Didcot OX11 0DE, UK

^6^ Medical, Molecular and Forensic Sciences, Murdoch University, Murdoch 6150, Australia

**Supporting Information**

**Supplementary Methods**

**Evaluation of compressive and torsional response of isolated intervertebral disc FEMs with elastic and viscoelastic material properties**

To estimate the impact of viscoelasticity on the normalised apparent axial compression modulus and torsional mechanics of the isolated intervertebral discs (n = 3, comprised of a nucleus pulposus and five-layer annulus fibrosus, **Figure S3a**) under quasi-static loading, additional simulations were conducted using the intervertebral disc FEMs with simplified boundary conditions. The superior and inferior surfaces of the intervertebral disc FEMs were identified based on anatomy, and their approximate geometric centres were used to define reference points: RP1 (superior surface) and RP2 (inferior surfaces). Kinematic coupling constraints were applied to tie all nodes on the superior surface to RP1, and all nodes on the inferior surface to RP2. For compressive modulus estimation, RP1 was encastred in all translational and rotational degrees of freedom (U1 – U3, UR1 – UR3 = 0). A uniaxial displacement of 40 μm was applied to RP2 in the Z direction at a displacement rate of 0.1 mm/min (taking 24 s to reach 40 μm displacement), followed by a 1200-second relaxation period with 40 μm displacement maintained.

For elastic models, material properties in **Table 3** were used for the nucleus pulposus and those in **Table 4** were used for the annulus fibrosus. For viscoelastic models, material properties in **Table S2** were used for the nucleus pulposus and the annulus fibrosus. To perform a geometry-based modulus normalisation, the geometric approximation of the intervertebral disc on the transverse plane is carried out using ellipses, from which the major and minor axes and areas of the annulus fibrosus [F_a_ (mm), F_b_ (mm), A_1_ (mm^2^)] and the nucleus pulposus [N_a_ (mm), N_b_ (mm), A_2_ (mm^2^)] were measured (**Figure S3b**). The height of the intervertebral disc was calculated using the average height of the anterior (h_1_, mm) and posterior (h_2_, mm) intervertebral disc (**Figure S3b**).

The reaction force-displacement curves at RP2 were generated to calculate the apparent compressive stiffness (S_comp_, N/mm) and the normalised apparent compressive modulus (E_comp_, MPa) using formula (1) and (2)[1].

*S_comp_ =* $\frac{F}{U}$	(1)

*E_comp_* = $\frac{S_{comp} \times h}{A}$	(2)

Where

*F* is the reaction force (N) at RF2.

*U* is the displacement (mm) of the boundary condition.

*h* is the height (mm) of the intervertebral disc.

*A* is the transverse area (mm^2^) of intervertebral disc, including both the annulus fibrosus (A_1_, mm^2^) and nucleus pulposus areas (A_2_, mm^2^).

For the evaluation of the torsional response, the same geometry and material properties were used. The boundary conditions were modified as follows. A rotational displacement of 0.1 radians (5.73°) about the Z-axis was applied to RP2 at a rate of 0.25 rad/min (taking 24 s to reach 0.1 radians displacement), followed by a 1200-second relaxation period. The polar moment of inertia (J, mm^4^) of the intervertebral disc was calculated as a hollow ellipse excluding the nucleus pulposus using formula (3)[1]. The moment-time curves at RP2 were generated to calculate the apparent torsional stiffness (K_tors_, N·mm/rad) and the normalised apparent torsional stiffness (G_tors_, MPa or MPa/°) using formula (4) and (5)[1].

*J =*$\frac{64 \times[F_{b}{F_{a}}^{3}+ {F_{b}}^{3}F_{a}-(N_{b}{N_{a}}^{3}+ {N_{b}}^{3}N_{a})]}{\pi}$	(3)

*K*_tors_ *=* $\frac{T}{\theta}$	(4)

*G_tors_* = $\frac{K_{tors} \times h}{J}$	(5)

Where

*F_a_* is the length of the major axis (mm) of the annulus fibrosus' elliptical approximation.

*F_b_* is the length of the minor axis (mm) of the annulus fibrosus' elliptical approximation.

*N_a_* is the length of the major axis (mm) of the nucleus pulposus' elliptical approximation.

*N_b_* is the length of the minor axis (mm) of the nucleus pulposus' elliptical approximation.

*T* is the reaction moment (N·mm) at RF2.

*θ* is the rotational displacement (rad) of the boundary condition.

*h* is the height (mm) of the intervertebral disc.

All simulations were performed using dynamic, implicit analysis step in Abaqus/Standard.

**FEM of the lumbar segments with elastic or viscoelastic intervertebral discs and comparison with the experiment**

Finite element models of the three rat lumbar segments (**Figure S4a**) with the above mentioned elastic or viscoelastic intervertebral discs were used to simulate a single loading step 1 (40 μm displacement), then compared with the experimental measurement acquired from the Deben CT5000 stage. Apart from the intervertebral disc, the other parts of the models and the modelling methods remained identical to those previously described in the ‘FEM of the rat lumbar segment’ in the Experimental Section unless otherwise stated. A loading rate of 0.1 mm/min (taking 24 s to reach 40 μm displacement) was applied, followed by a 1200-second relaxation. All simulations were performed using dynamic, implicit analysis step in Abaqus/Standard. Reaction force-time curves were extracted from the history output for analyses.

**Prediction of VEP mechanics under different motions**

To demonstrate a potential application of the optimised FEM, we used FEM to simulate different motions and evaluated the resultant strains in VEPs. Based on the FEM generated from the preloading step (1N, equivalent to ∼40 % body weight; to provide a baseline for quadrupedal posture), a series of boundary conditions for simulating different motions were applied to the rat lumbar segment FEM, including 13° of flexion, 2° of extension, 6° of lateral flexion, and 2° of rotation, respectively (**Figure S5**). All boundary conditions were applied to reference points RP1 and RP2 similar to the aforementioned in Methodology, and the displacements have constrained degrees of freedom except the direction of motion. Displacements were applied to RP1 while RP2 was encastred. The shear strains on the cranial and caudal VEPs were then calculated and visualised in the implicit solver.

**Sensitivity analysis on material and structural modifications**

Model with reduced moduli and geometric density was created in Avizo 3D using 3D erosion in the segmentation module. In the segmentation module, vertebral volume was selected, followed by the application of the built-in selection shrinkage algorithm. A total of 15 times of shrinkage were applied to enlarge the pores and thin the trabecula of the vertebrae. This allows an estimate of decreased trabecular volume in the model by ~30%[2]. After that, the erosive vertebral volume was converted into surface data with a simplified minimum edge length of 32.5 μm. The tetrahedral mesh was then created based on the surface data and exported as INP files. The vertebrae INP file was imported into Abaqus CAE to replace the original parts, and the elastic modulus decreased by 67% (3.96 GPa for cortical bone and 49.5 MPa for cancellous bone) in the vertebrae, and decreased by 33% (86.4 MPa) in the VEPs[2]. Regarding the model with increased moduli, the model retained vertebral and VEP trabecular microstructure[3], but the elastic modulus was increased in the vertebrae and VEP parts [4-5]. The elastic modulus was increased by 50% in the vertebrae (18.0 GPa for cortical bone and 225.0 MPa for cancellous bone) and by 30% in the VEPs (167.6 MPa). Simulation of uniaxial compression was performed in Abaqus by applying boundary condition of 3 N concentrated force (along Z direction with constrained zero force at X and Y directions) at RP1 while RP2 was encastred. The force applied is equivalent to 100% body weight of a rat. Model integration, interactions, job analyses, and visualisation were identically performed as described in the Experimental Section in the main text. The first principal stress, first principal strain, and third principal strain were obtained for statistical and visual analyses.

**Statistical analysis**

All statistical analyses were performed using SPSS Statistics and plotted using Origin. For the three isolated intervertebral disc FEMs, paired t-tests were conducted at each time point to compare the modulus values between elastic and viscoelastic models derived from the same disc (n = 3). To compare the differences in peak and relaxed forces among the experiment, elastic and viscoelastic lumbar segment models, Levene’s test was performed to assess the homogeneity of variances. In this case, variances were equal (p > 0.05), thus standard one-way ANOVA was performed. Effect sizes were calculated using eta-squared (η²), epsilon-squared, and omega-squared, with η² interpreted as small (1%), medium (6%), and large (14%) effects. For post hoc comparisons, LSD tests were used under the assumption of equal variances, and Tamhane’s T2 test was applied when variances were unequal. Where applicable, data are presented as mean ± standard deviation, with 95% confidence intervals. A significance level of p < 0.05 was considered statistically significant.

**Supplementary Note 1: Rationality of applying human lumbar material properties on rat lumbar FEMs**

The geometric comparison between human and rat lumbar reveals both similarities and differences that affect the rationality of cross-species material property applications. Although rat vertebrae exhibit a slenderer geometry compared to human vertebrae, the width-to-depth axial aspect ratios demonstrate similarity between species (**Figure S6**)[6]. This geometric conservation suggests that the fundamental axial and shear loading patterns and stress distributions in the axial direction are comparable between rat and human lumbar segments[6-7]. However, the slenderer rat vertebrae may introduce variations to sagittal or lateral bending loads.

In validation studies on the mechanical similarity between rat and human discs, it is demonstrated that the apparent disc mechanics is comparable after geometric normalisation. Elliott et al.[8] exhibited a compressive modulus of 2 – 4 MPa and a torsional modulus of 5 – 11 MPa for rat lumbar discs, well compared to a compressive modulus of 3 – 9 MPa and a torsional modulus of 2 – 9 MPa for human lumbar discs (**Table S3** and **S4**). Similarly, Orías et al.[9] reported a torsional stiffness of 0.081 ± 0.026 MPa/° for rat lumbar discs (axially loaded), fell within the range of 0.024 – 0.21 MPa/° for human lumbar discs (**Table S5**). Moreover, Showalter et al.[10] tested the torsional stiffness of calf, pig, baboon, goat, sheep, rabbit, rat, and mouse lumbar and compared to human lumbar. The results show that after geometric normalisation, the torsional stiffness of all animal lumbar discs have no significant difference from human (p > 0.05), except sheep and pig. The collagen content in the annulus discs also show similarity across species[10]. In our isolated rat disc FEMs (n = 3), despite the use of human material properties or elasticity/viscoelasticity, the instantaneous and relaxed normalised apparent compressive modulus, torsional modulus, and torsional stiffness are still within the reported range of rat or human lumbar disc mechanics (**Figure S3c–S3e**, **Table S3–S5**). These conserved structural features support the assumption that disc mechanics are governed by similar material principles across mammalian species.

Due to the limited availability of individual components of rat spine in literature, we applied human properties on the rat models as a compromise and to avoid unevidenced scaling or assumption. Above findings support that this method is appropriate and still can provide meaningful insights. However, several limitations should be acknowledged. First, the current rat-human comparability assumption is primarily established based on compressive and torsional responses, without involving bending loads. Second, the comparability is mainly validated at the organ level but not at the fibre level. Although macroscopic mechanical responses show similarity, differences in fibre arrangement, inter-lamellar connections, and the proportion of collagen types may introduce variations at finer scales. Nevertheless, given that this study primarily focuses on organ-level mechanical behaviour, the potential impact of these microscale differences on overall results may be relatively limited.

**Supplementary Note 2: Evaluation of impact of viscoelasticity in quasi-static experiments**

We performed comparative analyses on both isolated intervertebral disc and lumbar segment models. The results show that the viscoelastic FEMs exhibit time-dependent mechanical behaviour under quasi-static loading conditions (0.1 mm/min loading rate with 1200-second relaxation).

For isolated intervertebral disc models, the peak normalised apparent compressive modulus and torsional modulus of the viscoelastic models are 184% ± 11% (p < 0.001) and 202% ± 10% (p < 0.001) higher than those of elastic models at the completion of the loading phase, respectively (time = 24 s) (**Figure S3c–S3e**, **Table S3–S5**). In the following 1200-second relaxation stage, both the compressive and torsional moduli of the viscoelastic models gradually reduce, approaching long-term elastic response. At time = ~600 s onward, it shows no statistical significance (p > 0.05) between elastic and viscoelastic models for both moduli. By the end of relaxation (time = 1224 s), the normalised apparent compressive modulus and torsional modulus of the viscoelastic models are 7% ± 4% (p > 0.05) and 8% ± 5% (p > 0.05) lower than those of elastic models, respectively (**Figure S3c–S3e**, **Table S3–S5**).

For lumbar segment models, **Figure S4b** shows representative total reaction force curves from the experimental measurements and the models with either an elastic or viscoelastic intervertebral disc. In statistics, the peak total reaction force of the viscoelastic models is 320% ± 45% (p < 0.01, η² = 78.8%) higher than that of the elastic models. Even after relaxation, the relaxed total reaction force of the viscoelastic models is 203% ± 35% (p < 0.05, η² = 64.1%) higher than that of the elastic models. The total reaction force at the completion of relaxation of the elastic models is relatively closer to the experimental measurement than the viscoelastic models (**Figure S4c**), nevertheless, both the viscoelastic and elastic models show a good agreement (p > 0.05) with the experimental measurements at the completions of loading and relaxation stages.

In both isolated intervertebral disc and lumbar segment models, the greatest difference between elastic and viscoelastic models is found at the completion of the loading stage. The time-dependent convergence of mechanical responses between elastic and viscoelastic models is observed during the relaxation phase. These reflect the stress relaxation characteristics inherent to viscoelastic materials. The convergence of mechanical responses under quasi-static conditions can be attributed to the low loading rate of 0.1 mm/min and the extended relaxation period of 1200 s. Our loading rate is 1 – 2 orders of magnitude lower than typical biomechanical experiments[11-13], weakening the impact of viscoelastic effect. The extended relaxation time allows the viscoelastic models to approach their equilibrium state and shows no statistical significance (p > 0.05) to the elastic models from ~600 s onward.

In isolated intervertebral disc models, the differences between elastic and viscoelastic models become negligible after sufficient relaxation time, consistent with the stress relaxation behaviour of individual disc components. However, it is noted that in complete lumbar segment models, differences with statistical significance (p < 0.05, η² = 64.1%; or p < 0.01, η² = 78.8%) persisted between elastic and viscoelastic models even after relaxation. This discrepancy arises because in the loading phase, compared to the models with elastic intervertebral disc, the viscoelastic models exhibit higher instantaneous modulus, thus greater deformation in other elastic parts (vertebrae, growth plates, and vertebral/cartilaginous endplates). As the viscoelastic intervertebral disc undergoes stress relaxation, the elastic parts remain deformed and maintain high residual reaction.

These findings demonstrate that under our quasi-static experimental conditions, the mechanical behaviour of intervertebral discs can be effectively approximated using anisotropic elastic material properties (engineering constants). This equivalence simplifies computational analysis while maintaining physiological relevance. However, this simplification is strictly limited to the specific experimental setup and should not be generalised to cyclic, or high strain rate loading scenarios, where viscoelastic effects remain substantial and cannot be neglected.

**Supplementary Note 3: Example applications of the optimised FEM in biomechanics**

We exhibit two primary applications of this optimised FEM: revealing the key mechanical behaviours of VEP macrostructures under different motions and diseased conditions. We emphasise that this is intended solely to demonstrate potential applications. Thus, no biological conclusions have been drawn, nor has statistical significance been established.

In the simulation of flexion and extension along different axes, FEM showed that the sagittal and lateral protrusions retained most deformation even though their range varied under the different motions. Thus, sagittal-plane flexion and extension generated greater shear strain in sagittal than in lateral VEP protrusions (e.g., cranial VEP, a mean E23 shear strain of 0.5% on the sagittal, compared to 0.15% and 0.01% on the lateral and body, respectively), whilst lateral-plane flexion and rotation, yielded greater shear strain on the horizontal plane of the lateral VEP protrusions (e.g., cranial VEP, a mean E12 shear strain of 0.07% on the lateral, compared to -0.02% and 0.02% on the sagittal and body, respectively) (**Figure S5**).

As a second example, we used our optimised FEM to explore how artificial changes in material properties and geometry might influence strain distributions in the VEP. Specifically, we conducted a sensitivity analysis by adjusting the elastic modulus and geometric density of the vertebrae and VEP (**Figure S7a**). Under identical 3N load (equivalent to 100% rat body weight), we observed increased VEP strains in the model with reduced moduli and geometric density than in other conditions, particularly in the four VEP protrusions (**Figure S7b** and **7c**). Regional differences were also observed. For example, when compared to the baseline FEM, the lateral protrusions of cranial VEP experienced the maximum increase of mean third principal strain by 27%, whilst ~20% in the VEP body and the sagittal. The caudal VEP, in contrast, the minimum increase of mean third principal strain by only 3% was observed in the lateral protrusions, compared to ~12% in the VEP body and the sagittal. In the model with increased modulus, compressive strains evenly reduced over all regions of both cranial and caudal VEP, compared to the baseline FEM (by about 18 -20%) (**Figure S7b** and **S7c**).

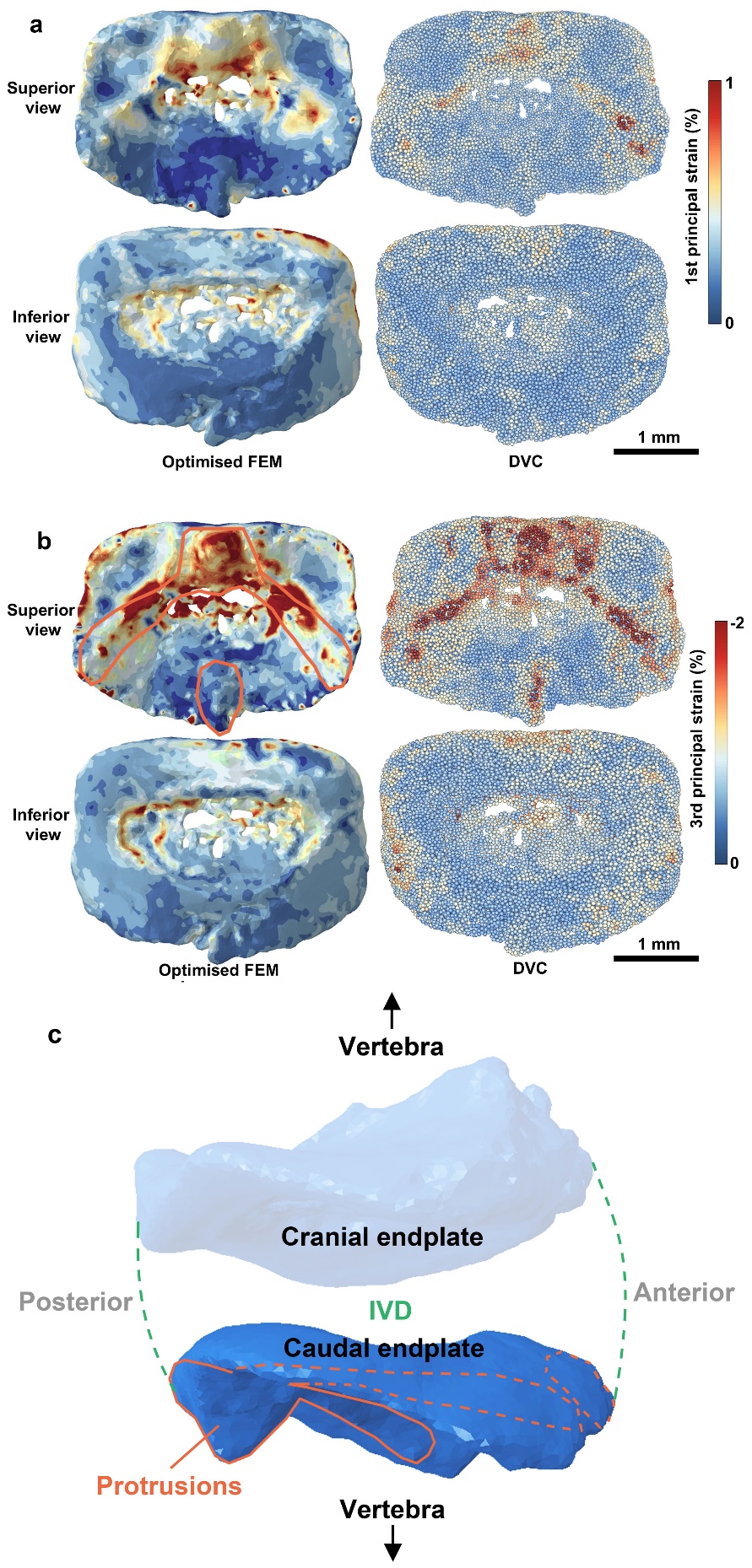

**Figure S1**. Visualisation of **(a)** first principal strain and **(b)** third principal strain on the caudal VEP optimised FEM and DVC point cloud reference under 80 μm compressive displacement. Orange lines indicate the protrusions of caudal VEP. (**c**) Schematic of the spatial relationship between the caudal VEP protrusions and other tissues. Orange dashed lines indicate the protrusions of caudal VEP.

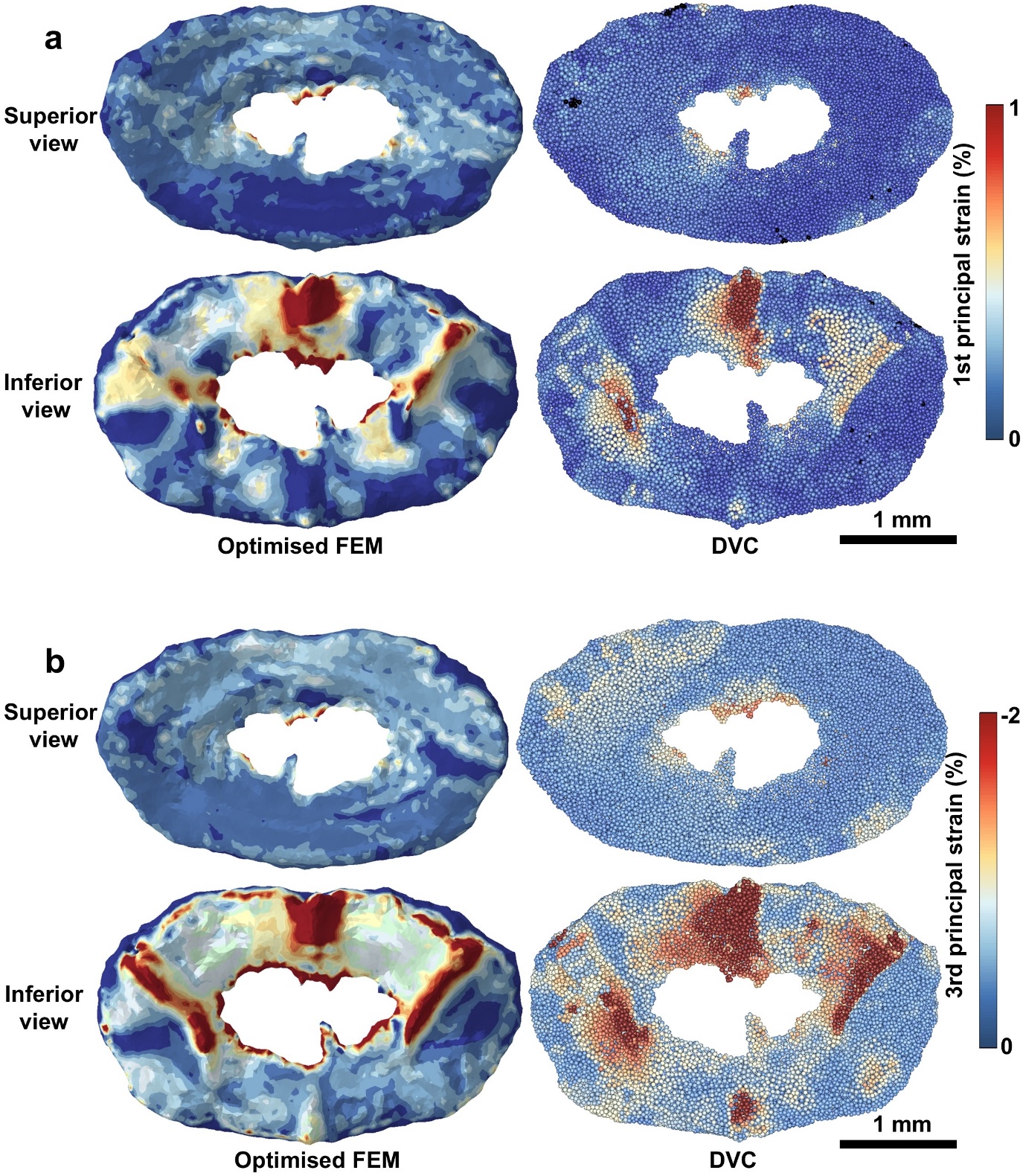

**Figure S2**. Visualisation of **(a)** first principal strain and **(b)** third principal strain on the cranial VEP optimised FEM and DVC point cloud reference at the second loading step (80 μm compressive displacement).

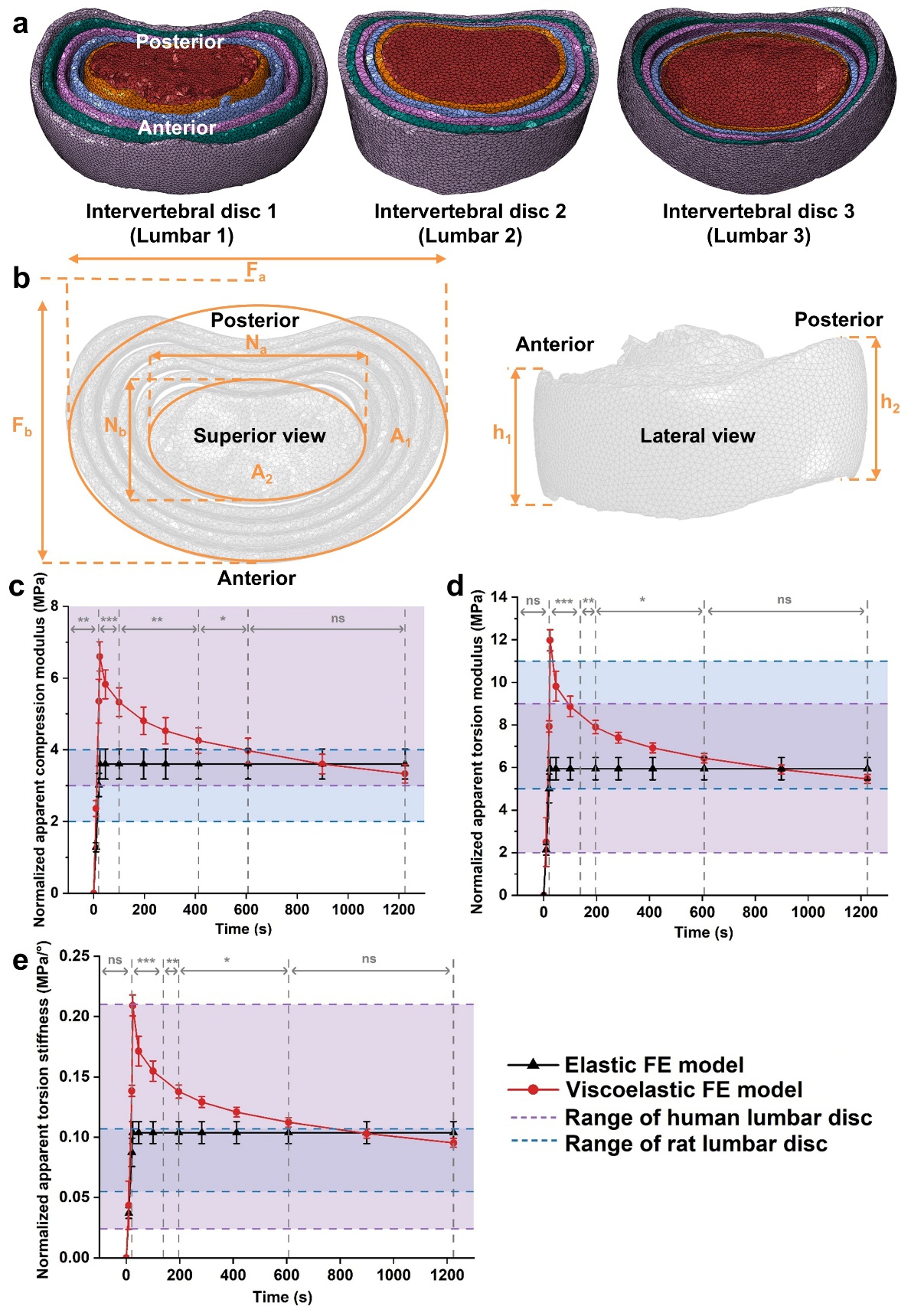

**Figure S3. Comparison between isolated elastic and viscoelastic rat intervertebral disc FEMs.** **(a)** Finite element models of three isolated rat intervertebral discs; **(b)** Geometric measurements of the intervertebral discs; **(c)** Normalised apparent compressive modulus-time curves under 40 μm uniaxial compression; **(d)** Normalised apparent torsional modulus-time curves under 0.1 radian (5.73°) rotation about the z-axis, and **(e)** conversion to torsional stiffness. The purple region indicates the reported range of human lumbar disc properties from the literature, while the blue region represents the reported range of rat lumbar disc properties. Not significant (ns): P > 0.05; *: P < 0.05; **: P < 0.01; ***: P < 0.001.

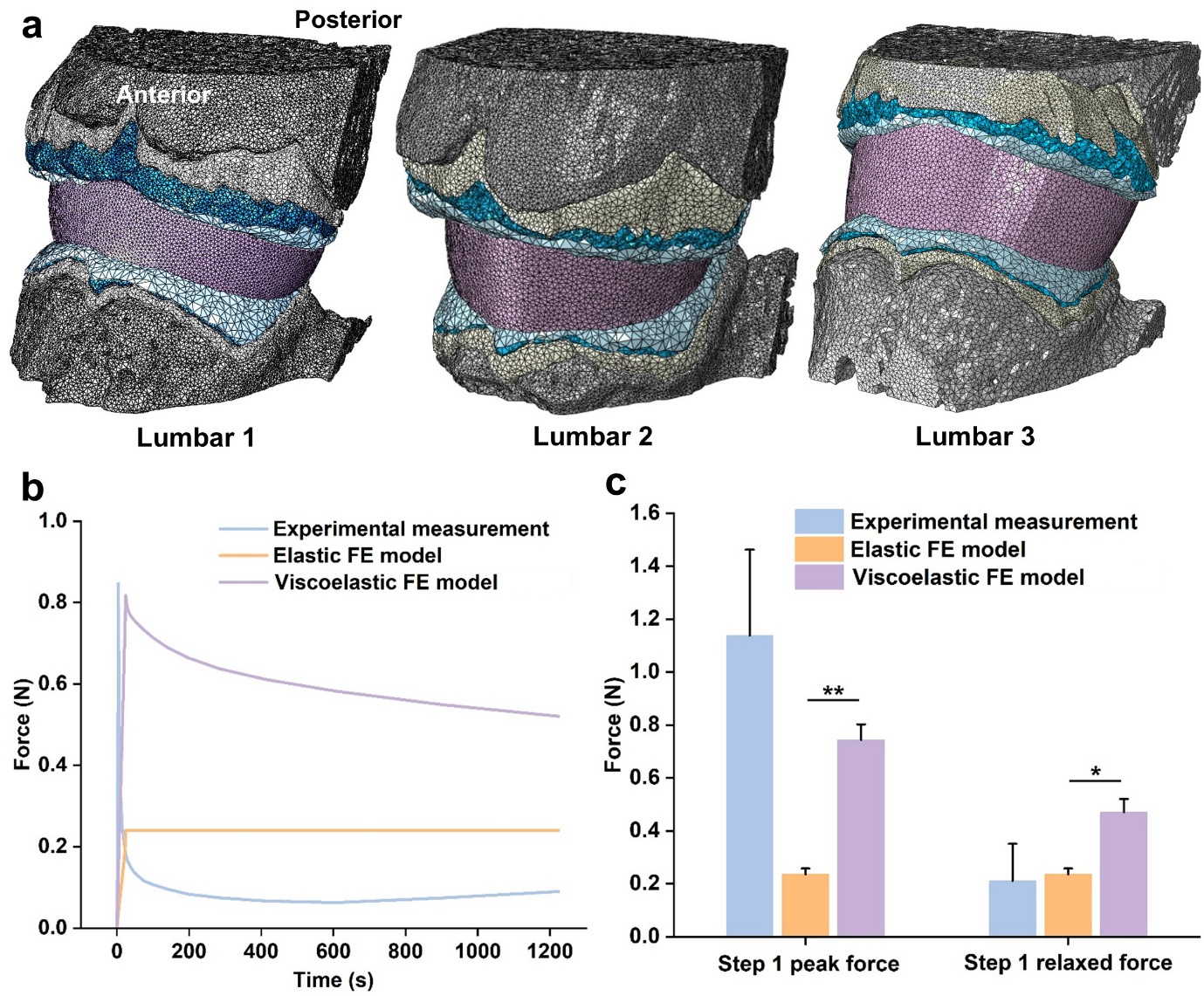

**Figure S4. Comparison of peak and relaxed forces between experiment and rat lumbar segment FEMs with elastic/viscoelastic intervertebral discs.** **(a)** Three rat lumbar segment FEMs with elastic/viscoelastic intervertebral discs used for the comparison; **(b)** Representative force-displacement curves of Lumbar 1, generated from experimental measurements and elastic//viscoelastic FEMs (loading step 1, 40 μm displacement); **(c)** Statistical analysis of peak and relaxed forces from experimental measurements and elastic//viscoelastic FEMs (loading step 1, 40 μm remote displacement). ns: P > 0.05; *: P < 0.05; **: P < 0.01; ***: P < 0.001. Compared to the experimental measurement.

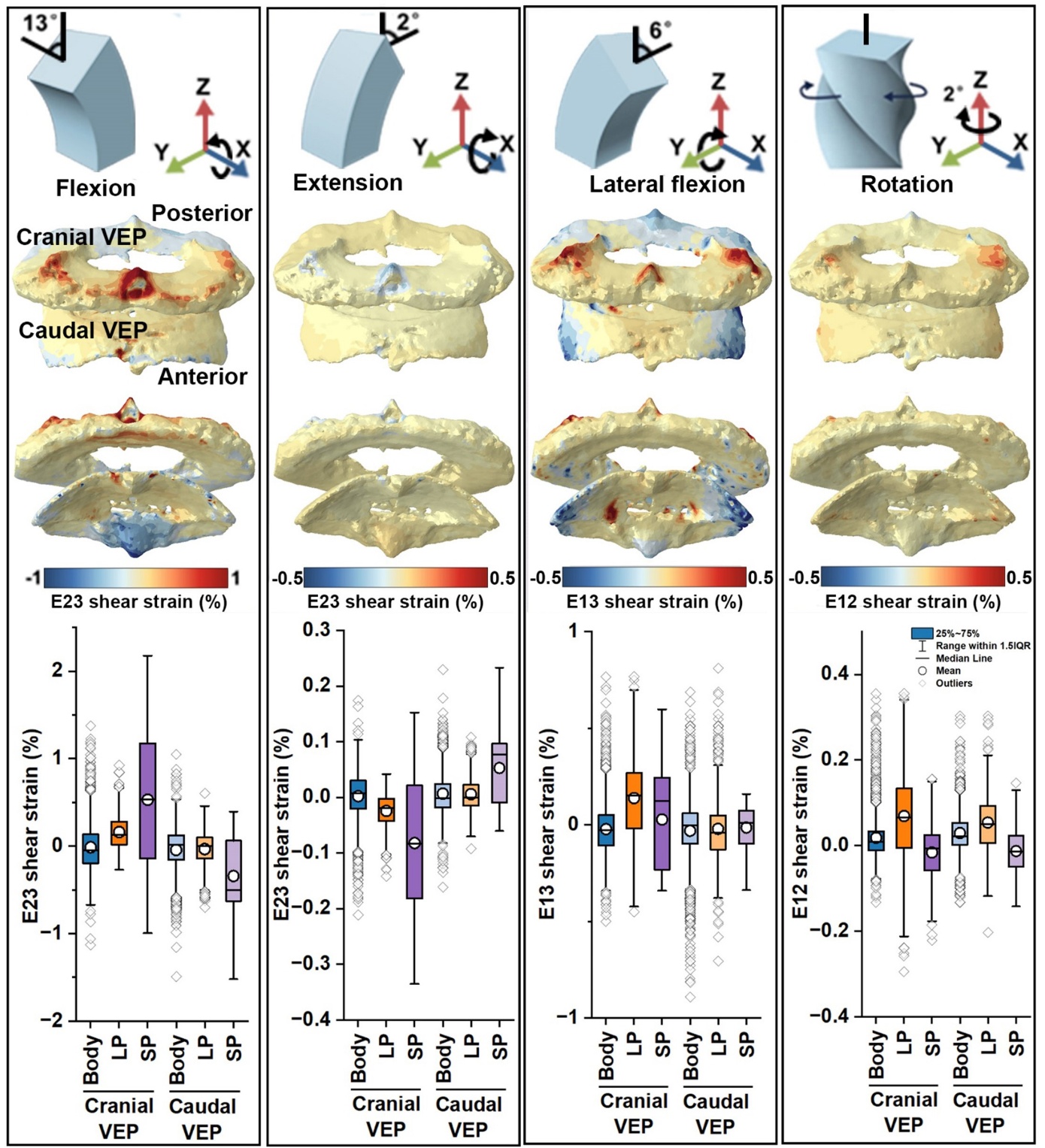

**Figure S5**. **Biomechanics of VEP macrostructures under flexion, extension, and rotation.** Shear strain distributions on sagittal (E23), coronal (E13), and horizontal (E12) planes of cranial and caudal VEPs under different motions. The box plots illustrate the strain distribution across different regions of the VEP surface on the vertebral side. Body: VEP body; LP: Lateral protrusions; SP: Sagittal protrusions.

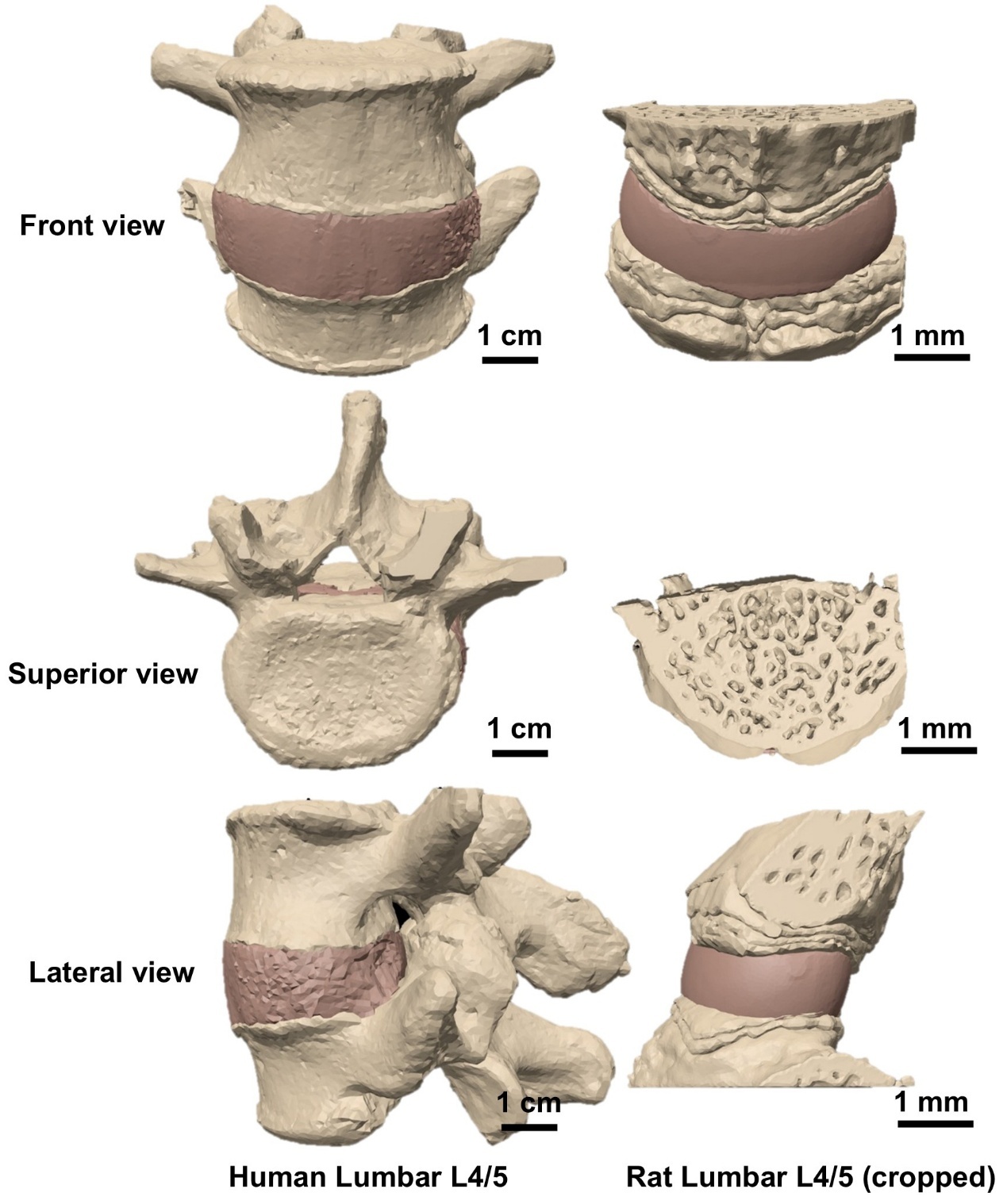

**Figure S6. 3D render of human and rat lumbar (cropped) L4/5 segments.**

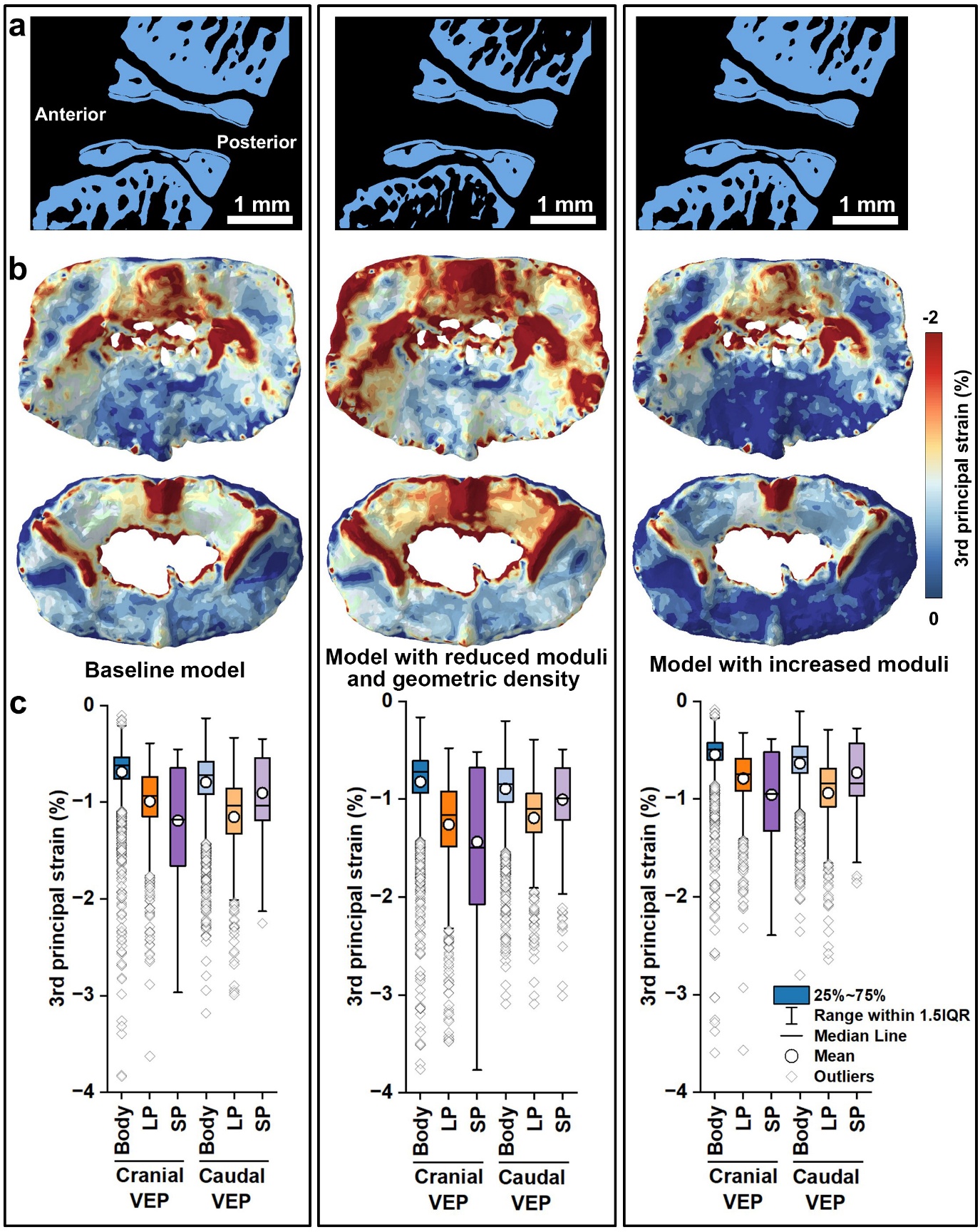

**Figure S7. Simulation and comparison of baseline model and models with adjusted moduli and geometric density.** **(a)** Orthoslice of lumbar FEMs under different conditions; **(b)** Visualisation of third principal strain on the cranial and caudal VEPs; **(c)** Box plots of third principal strain distribution across different regions of the VEP surface on the vertebral side. Body: VEP body; LP: Lateral protrusions; SP: Sagittal protrusions.

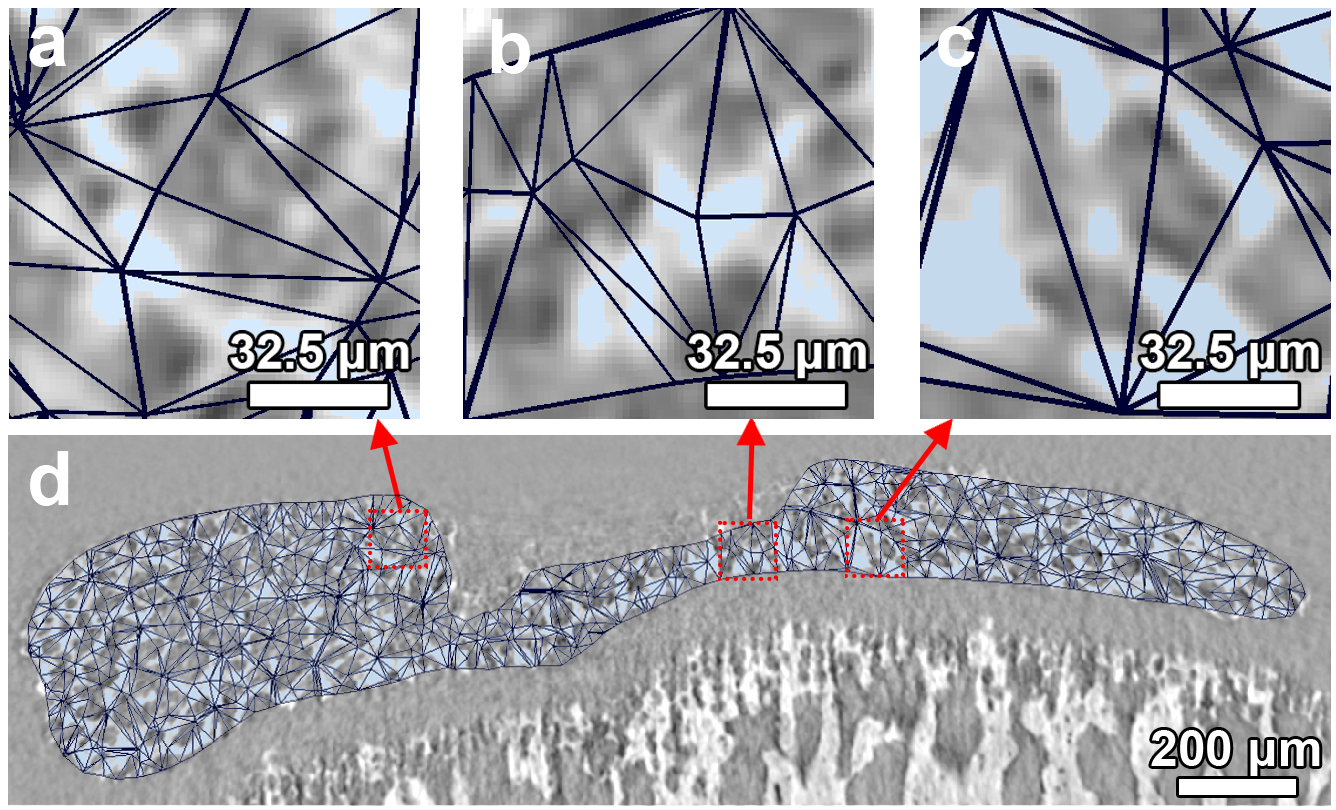

**Figure S8**. 2D planar slice of FEM mesh (black triangle) with superimposed sCT image of VEP in grayscale. **(a–c)** Representative zoom-in views show that each mesh element incorporates both bony VEP (high intensity) and unmineralised components (medium-low intensity). The minimum edge length of elements is 32.5 μm; (d) Overview of FEM mesh for VEP.

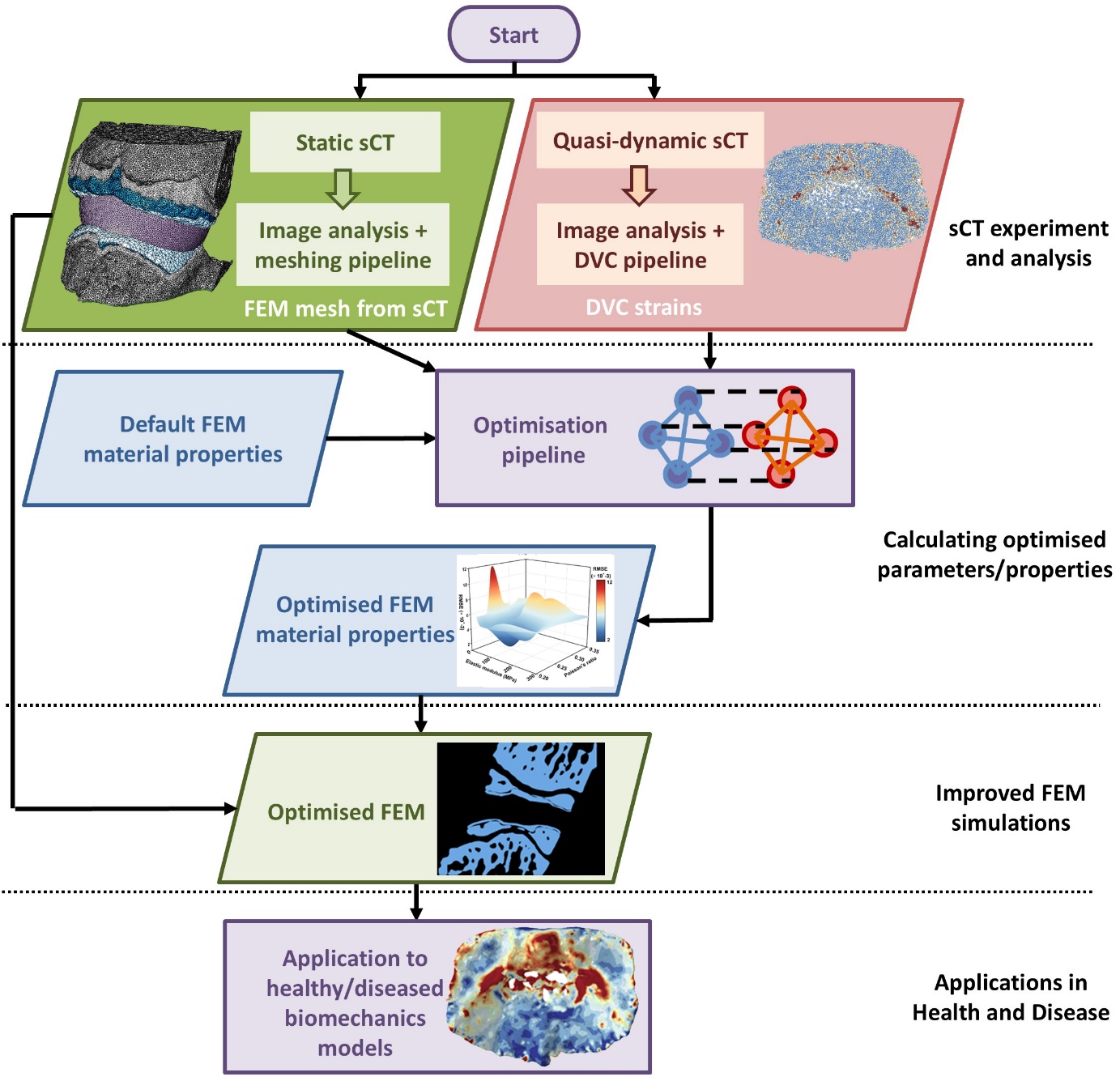

**Figure S9. Workflow of the methods.**

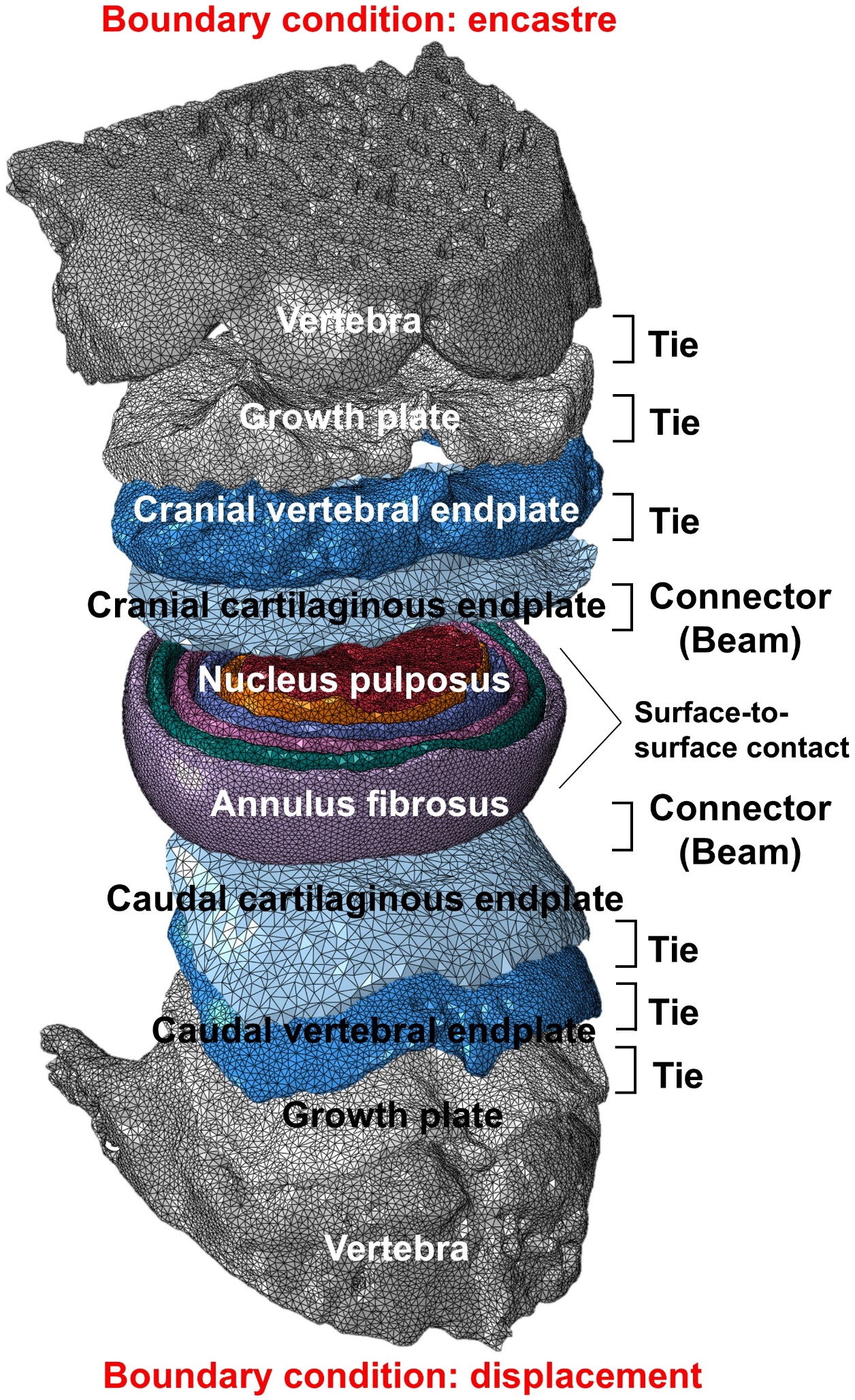

**Figure S10. Interaction and boundary conditions between FEM parts.**

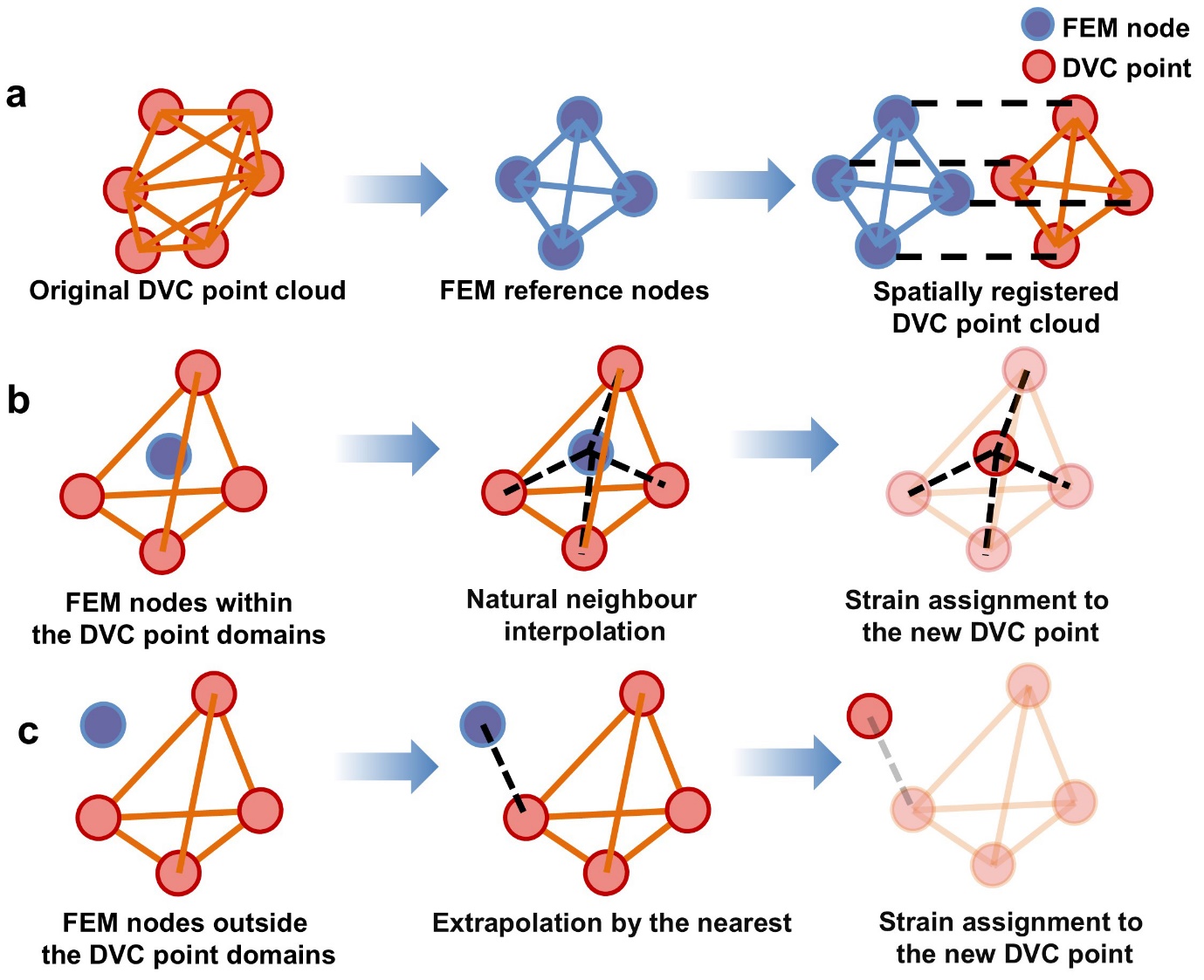

**Figure S11. Spatial registration of the DVC point cloud to FEM reference nodes.** **(a)** spatial registration of the DVC point cloud; **(b)** Natural neighbour interpolation method for new FEM-referenced DVC points within the original DVC point domains; **(c)** Extrapolation by the nearest method for new FEM-referenced DVC points outside the original DVC point domains.

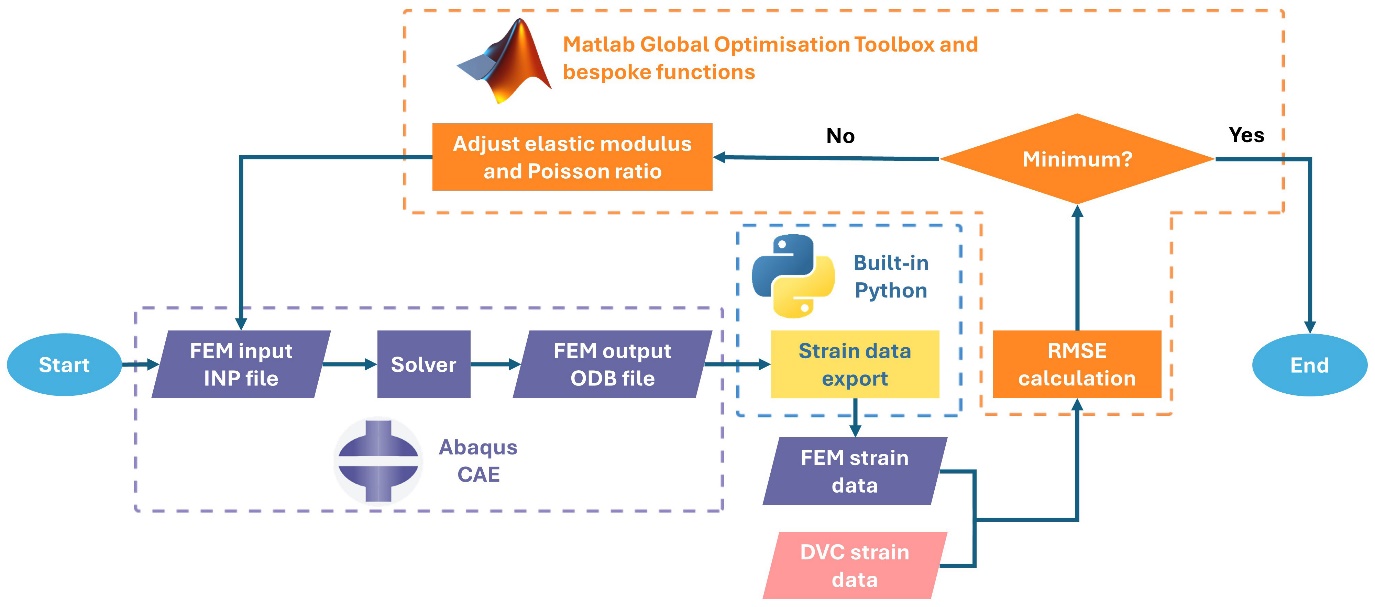

**Figure S12. Flow chart of the inversion pipeline.**

**Table S1.** Statistics of percentage difference in mean values of first principal strain between DVC measurements and FEM predictions at different compression steps.

| **Samples** | **Type of strain** | **Compression steps** | **Percentage difference in mean values of first principal strain (DVC vs FEM, %)** |
| --- | --- | --- | --- |
| Caudal VEP | First principal strain | 1 | 7.5 |
|  |  | 2 | 7.2 |
|  |  | 3 | 8.3 |
|  | Third principal strain | 1 | 5.2 |
|  |  | 2 | 1.1 |
|  |  | 3 | 3.9 |
| Cranial VEP | First principal strain | 1 | 6.4 |
|  |  | 2 | 7.9 |
|  |  | 3 | 1.2 |
|  | Third principal strain | 1 | 1.5 |
|  |  | 2 | 3.7 |
|  |  | 3 | 7.2 |

| **Materials** | **Instantaneous elastic modulus** | **Viscoelasticity (Prony series)[14-15]** | | | **Element type** |
| --- | --- | --- | --- | --- | --- |
|  |  | **Relaxation of** | | |  |
|  |  | **Shear** | **Bulk** | **Time (s)** |  |
| Nucleus pulposus | Isotropic elastic: modulus = 1 MPa, Poisson's ratio = 0.4999 (imconpressible)[16] | 0.6375 | 0 | 0.141 | C3D4H |
|  |  | 0.1558 | 0 | 2.21 |  |
|  |  | 0.1202 | 0 | 39.9 |  |
|  |  | 0.0383 | 0 | 266 |  |
|  |  | 0 | 0.8000 | 500 |  |
| Annulus fibrosus | Anisotropic Hyperelastic (Holzapfel-Gasser-Ogden)^*^: c10 = 0.85, d =0.0001, k1 = 2.8, k2 = 90, kappa = 0[17] | 0.3392^**^ | 0.3392^**^ | 3.45 | C3D4 |
|  |  | 0.0000^**^ | 0.2550^**^ | 100 |  |
|  |  | 0.3064^**^ | 0.1267^**^ | 1000 |  |
|  |  | 0.0920^**^ | 0.1275^**^ | 5000 |  |

**Table S2.** Material properties of the isolated intervertebral disc FEMs according to the references with minor modifications.

Note: *The strain energy potential-based material properties for an anisotropic, fibre-reinforced annulus fibre-matrix continuum; **A factor of 0.85 was applied to the reported Prony series for the annulus matrix in [14], as the matrix is the primary contributor to the viscoelastic behaviour and accounts for ~85% of the volume in the annulus fibrosus[15].

| **Sample reference** | **Material type of disc** | **Compressive modulus (MPa)** | | | | **Reported range (rat, MPa)[1]** | **Reported range**  **(human, MPa)[1]** |
| --- | --- | --- | --- | --- | --- | --- | --- |
|  |  | **Peak** | **Statistics** | **Relaxed** | **Statistics** |  |  |
| 1 | Elastic | 3.97 | 3.61 ± 0.42 | 3.97 | 3.61 ± 0.42 | 2 – 4 | 3 – 9 |
| 2 |  | 3.02 |  | 3.02 |  |  |  |
| 3 |  | 3.83 |  | 3.83 |  |  |  |
| 1 | Viscoelastic | 6.97 | 6.60 ± 0.41 | 3.60 | 3.33 ± 0.26 |  |  |
| 2 |  | 6.04 |  | 2.98 |  |  |  |
| 3 |  | 6.80 |  | 3.41 |  |  |  |

**Table S3.** Normalised apparent compressive modulus of the isolated intervertebral disc FEMs.

| **Sample reference** | **Material type of disc** | **Torsional modulus (MPa)** | | | | | **Reported range (rat, MPa)[1]** | **Reported range**  **(human, MPa)[1]** |
| --- | --- | --- | --- | --- | --- | --- | --- | --- |
|  |  | **Peak** | **Statistics** | **Relaxed** | | **Statistics** |  |  |
| 1 | Elastic | 6.04 | 5.95 ± 0.52 | 6.04 | 5.95 ± 0.52 | | 5 – 11 | 2 – 9 |
| 2 |  | 6.54 |  | 6.54 |  |  |  |  |
| 3 |  | 5.27 |  | 5.27 |  |  |  |  |
| 1 | Viscoelastic | 11.96 | 11.98 ± 0.49 | 5.45 | 5.46 ± 0.21 | |  |  |
| 2 |  | 12.60 |  | 5.72 |  |  |  |  |
| 3 |  | 11.39 |  | 5.21 |  |  |  |  |

**Table S4.** Normalised apparent torsional modulus of the isolated intervertebral disc FEMs.

| **Sample reference** | **Material type of disc** | **Torsional stiffness (MPa/°)** | | | | **Reported range (rat, MPa/°)[18]** | **Reported range**  **(human, MPa/°)[18]** |
| --- | --- | --- | --- | --- | --- | --- | --- |
|  |  | **Peak** | **Statistics** | **Relaxed** | **Statistics** |  |  |
| 1 | Elastic | 0.105 | 0.104 ± 0.009 | 0.105 | 0.104 ± 0.009 | 0.081 ± 0.026 (axially loaded);  0.091±0.033 (unloaded) | 0.024 – 0.210 |
| 2 |  | 0.114 |  | 0.114 |  |  |  |
| 3 |  | 0.092 |  | 0.092 |  |  |  |
| 1 | Viscoelastic | 0.209 | 0.209 ± 0.09 | 0.095 | 0.095 ± 0.004 |  |  |
| 2 |  | 0.220 |  | 0.100 |  |  |  |
| 3 |  | 0.199 |  | 0.091 |  |  |  |

**Table S5.** Normalised apparent torsional stiffness of the isolated intervertebral disc FEMs.

**Table S6.** The reported elastic modulus and Poisson's ratio of endplates in the literature.

| **Testing method** | **Type of endplate** | **Elastic modulus (MPa)** | **Poisson's ratio** | **Species** | **Spine segments** | **References** |
| --- | --- | --- | --- | --- | --- | --- |
| Indentation | Bony | 6,000 | 0.3 | Human | L3-L4 | [19] |
|  | Bony | 1,000 | 0.4 | Human | T11-L1 | [20] |
|  | Bony | 245 | / | Human | L1-L2 | [21] |
|  | Bony and cartilaginous | 220 | / | Rabbit | L1-L7 | [22] |
| Universal testing machine | Bony and cartilaginous | 600 | 0.3 | Human | C4-C7 | [23] |
|  | Bony and cartilaginous | 600 | 0.3 | Human | C1-T1 | [24] |
|  | Bony and cartilaginous | 500 | 0.4 | Human | C0–C7 | [25] |
|  | Bony and cartilaginous | 500 | 0.4 | Human | C2-C7 | [26] |
|  | Bony and cartilaginous | 500 | 0.25 | Human | L1-L5 | [27] |
|  | Bony and cartilaginous | 23.8 | 0.4 | Human | L1-S1 | [28] |
| Unspecified | Bony and cartilaginous | 1,200 | 0.29 | Human | L3-S1 | [29] |
|  | Bony and cartilaginous | 1,200 | 0.29 | Human | L1-L5 | [30] |
|  | Bony | 1,000 | 0.3 | Human | T11-L5 | [31] |
|  | Bony and cartilaginous | 1,000 | 0.3 | Human | L4-L5 | [16] |
|  | Bony and cartilaginous | 500 | 0.25 | Human | L1-L5 | [32] |
|  | Bony and cartilaginous | 500 | 0.4 | Human | C3-C7 | [33] |
|  | Bony | 25 | 0.25 | Human | L3-L4 | [34] |
|  | Bony and cartilaginous | 25 | 0.3 | Human | L4-S1 | [35] |
|  | Bony | 24 | 0.4 | Human | L4-L5 | [36] |
|  | Bony and cartilaginous | 24 | 0.4 | Human | L1-S1 | [37] |

**Table S7.** Material properties used in the sensitivity test.

| **Part** | **E_min_ (Isotropic, MPa)** | **E_max_** **(Isotropic, MPa)** | **Reference** |
| --- | --- | --- | --- |
| Vertebrae (Cortical bone) | 10000 | 12000 | **E_min_** [38], **E_max_** [39] |
| Vertebrae (Cancellous bone) | 100 | 450 | **E_min_** [40], **E_max_** [39] |
| Growth plates | 7.5 | 12 | **E_min_** [41], **E_max_** [42] |
| Cartilaginous endplates | 23.8 | 24 | **E_min_** [43], **E_max_** [44] |
| Nucleus pulposus | 0.2 | 1 | **E_min_** [45], **E_max_** [40] |
| Annulus fibrosus (Layer 1 – 5) | 175 | 750 | **E_min_** [46], **E_max_** [47] |
